# A *Slc5a6* Deficient Mouse Model Reveals a Metabolically Driven Dilated Cardiomyopathy with Therapeutic Potential for Vitamin-Based Intervention

**DOI:** 10.1101/2025.06.06.658273

**Authors:** Millie O. Fullerton, Lauren C. Phillips, Rachael E. Redgrave, Vincent Haufroid, George Merces, Scott T. Kerridge, Gavin D. Richardson, Nathalie Mercier, Dominique Roland, Joseph P. Dewulf, John Burn, Simon D. Bamforth, Helen M. Phillips

## Abstract

**Background and Aims:** The sodium-dependent multivitamin transporter, encoded by *SLC5A6*, mediates cellular uptake of the vitamins, biotin and pantothenic acid, both of which are essential cofactors for energy metabolism. Here, we report two families with *SLC5A6* mutations presenting with early-onset dilated cardiomyopathy (DCM). To investigate the link between vitamin deficiency and DCM, we generated a novel cardiac-specific *Slc5a6* knockout (*Slc5a6^cKO^*) mouse model and tested the therapeutic potential of vitamin supplementation.

**Methods:** Cardiac function in *Slc5a6^cKO^* mice was assessed by cardiac magnetic resonance imaging and ECG measurements. Histological, biochemical, and proteomic analyses were conducted to identify structural and metabolic changes. The impact of dietary biotin and pantothenic acid supplementation on disease progression was evaluated.

**Results:** *Slc5a6^cKO^* mice developed progressive cardiac dysfunction, manifesting as DCM with cardiac dilation, cardiomyocyte hypertrophy, fibrosis, impaired Coenzyme A synthesis, and metabolic imbalance, culminating in premature death by 26 weeks. Proteomic analysis revealed early mitochondrial metabolic disruption and extracellular matrix protein upregulation at 8 weeks, preceding overt cardiac dysfunction. Strikingly, vitamin supplementation from preconception onwards, prevented the cardiac phenotype, preserving cardiac structure, function, morphology and survival. This parallels the clinical outcome in one patient who received early vitamin treatment, compared to another who required a heart transplant following delayed vitamin treatment.

**Conclusions:** This study establishes a direct link between SLC5A6-mediated vitamin transport, mitochondrial function, and cardiac health. It highlights how vitamin deficiency contributes to DCM pathogenesis and supports early vitamin supplementation as a potential therapeutic strategy for metabolic cardiomyopathies.

**Translational perspective:** This study highlights the therapeutic potential of vitamin supplementation in treating dilated cardiomyopathy (DCM) caused by mitochondrial abnormalities. Using a cardiac-specific *Slc5a6* knockout mouse model, we demonstrated that deficiencies in key vitamins, biotin and pantothenic acid, impair mitochondrial energy metabolism, leading to DCM progression. Remarkably, vitamin supplementation preserved cardiac function, morphology, and survival, suggesting that restoring vitamin levels could be a promising therapeutic strategy for DCM and other cardiomyopathies linked to metabolic deficiencies. These findings could inform newborn screening programmes and clinical approaches for treating mitochondrial-related cardiac diseases by targeting specific vitamin deficiencies.

**Key Question:** What is the underlying molecular cause of early-onset dilated cardiomyopathy in patients with *SLC5A6* mutations, and can insights from a cardiac-specific knockout mouse model reveal potential metabolic mechanisms and therapeutic strategies involving vitamin supplementation?

**Key Finding:** Cardiac-specific deletion of *Slc5a6* in mice caused early mitochondrial dysfunction, metabolic derangement, and progressive dilated cardiomyopathy. Strikingly, early and continuous supplementation with biotin and pantothenic acid completely preserved cardiac structure, function, and survival, paralleling successful outcomes in patients treated early.

**Take Home Message:** This study establishes a mechanistic link between *SLC5A6* mutations, vitamin deficiency and mitochondrial abnormalities as a cause of dilated cardiomyopathy. Early vitamin supplementation prevents disease onset, highlighting the potential of targeted vitamin therapy in metabolic cardiomyopathies.

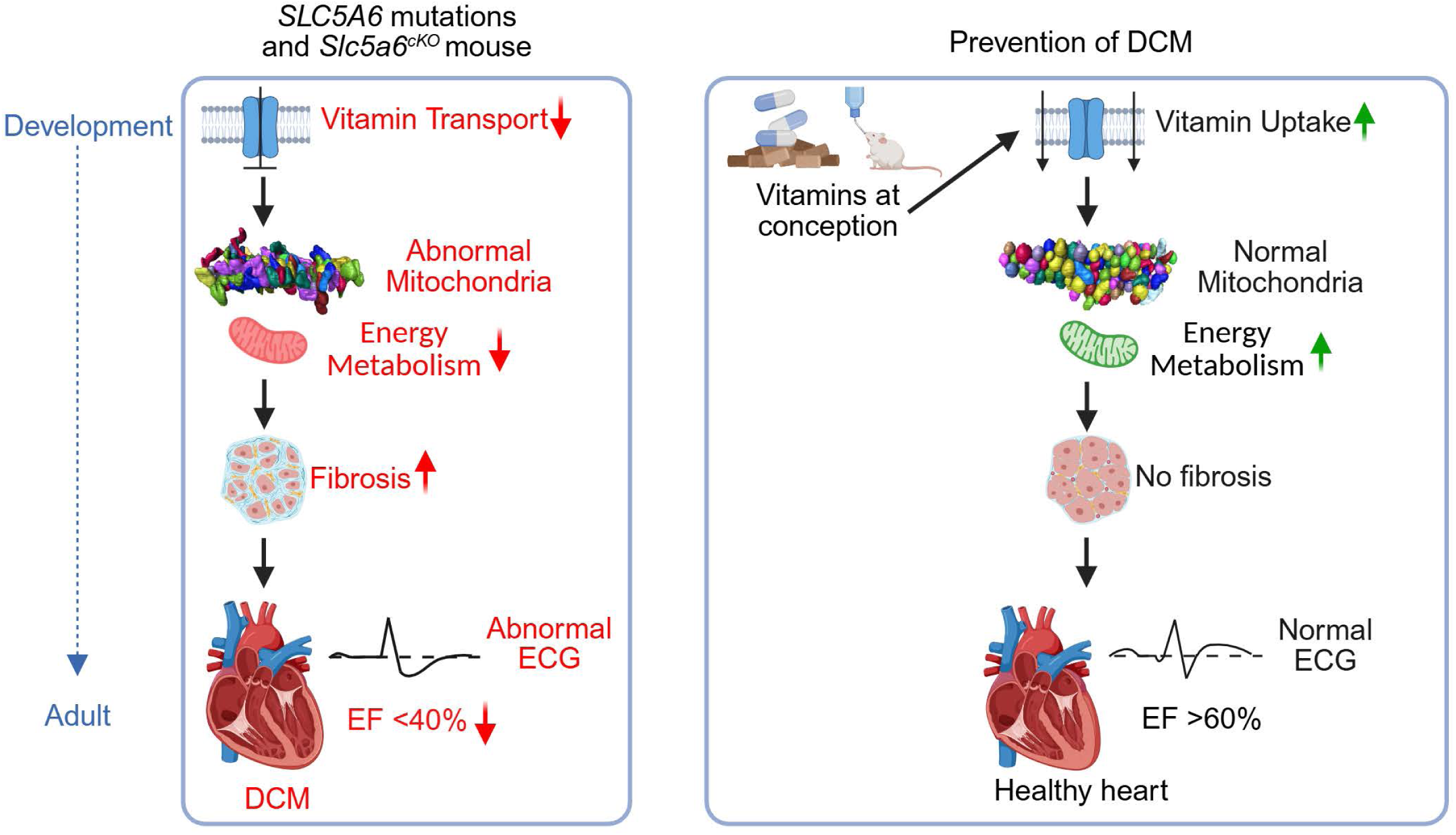

## Introduction

Dilated cardiomyopathy (DCM) is a leading cause of cardiovascular morbidity and mortality, affecting approximately 1 in 250 people in the UK, characterised by enlargement and dilation of one or more ventricles with impaired contractility, reduced left ventricular ejection fraction (EF) and heart failure. Paediatric cases of DCM are particularly severe, with over one-third of affected children requiring heart transplantation.^(1)^ The aetiology of DCM includes both genetic and environmental factors, such as mutations in sarcomeric genes^(2)^ or exposure to cardiotoxic drugs.^(3)^ Many cases of DCM, however, are idiopathic, and the complex pathological mechanisms underlying DCM are still not well understood. The heart is the most metabolically demanding organ in the body, requiring high ATP turnover, given the critical role of mitochondria in energy metabolism, disruptions in mitochondrial dynamics and function have been strongly linked to cardiovascular disease and heart failure.^(4)^

*SLC5A6* encodes the highly conserved and ubiquitously expressed sodium-dependent multivitamin transporter (SMVT), a member of the amino acid-polyamine organocation superfamily, sodium/solute symporters, which allow the unilateral transport of molecules across the cell membrane.^(5–7)^ It is required for the uptake of water-soluble vitamins biotin (vitamin B7), pantothenic acid (vitamin B5) and lipoic acid.^(5, 6, 8, 9)^ Biotin and pantothenic acid are acquired from the diet ^(5, 10)^ and are vital for energy production via the tricarboxylic acid (TCA) cycle, amino acid catabolism and fatty acid synthesis.

Patients with homozygous or compound heterozygous mutations in *SLC5A6* encompass an autosomal recessive disorder and exhibit a wide spectrum of multisystemic clinical manifestations, including developmental and growth delay, neurodegenerative disorders, gastrointestinal problems, immunodeficiency, seizures and osteopenia.^(11–23)^ However, the influence of *SLC5A6* mutations on cardiovascular health and function has not been studied.

Here, we describe three patients, two of whom are from a newly identified consanguineous family and compare it with a previously published patient^(18)^, both of which presented with early onset DCM and homozygous *SLC5A6* mutations. Family 1 has two affected siblings, one of whom died prematurely, while the other had a life-saving heart transplant, prior to receiving vitamin supplementation at a later stage. In the second family, the patient similarly presented with severe heart failure but had a successful response to early vitamin supplementation, thus preventing the need for a transplant.

To investigate an association between DCM and vitamin deficiency, we generated a novel cardiac-specific conditional deletion of *Slc5a6* in mice (*Slc5a6^cKO^*). These mice exhibited a progressive DCM-like phenotype, culminating in premature death around 26 weeks. We identified abnormalities in cardiac function and protein expression preceding the onset of histological manifestations. Metabolomic and proteomic analyses of *Slc5a6^cKO^*hearts revealed disruptions in metabolic pathways and an increase in fibrosis-associated proteins from as early as 8 weeks of age. Remarkably, vitamin supplementation completely ameliorated the DCM phenotype, with functional, pathological and protein expression changes completely absent.

## Methods

Detailed methods are provided in the Supplementary Material.

## Results

### Clinical manifestations and progression of DCM in two consanguineous families

In Family 1, which is of Pakistani ethnicity, three siblings were born to healthy consanguineous parents (Figure 1A). All pregnancies carried to full term without any complications. Sibling II-1 died at 24 months due to DCM. Patient II-2 developed an acute illness at 9 months, characterized by hypoglycaemia and lethargy, which rapidly progressed to respiratory failure, necessitating paediatric intensive care unit admission and mechanical ventilation for two weeks. During this time, total parenteral nutrition (TPN) was given, and clinical indicators of DCM were first identified. A subsequent admission occurred at 21 months due to cardiac dysfunction requiring inotropic support, and the patient developed pneumonia, left-sided paralysis, and seizures. Mild developmental delay was noted, and nutritional support was administered via nasogastric tube due to feeding difficulties. The patient then remained well for two years. However, at 4 years of age, the patient relapsed and experienced cardiac decompensation and had poor cardiac output, with imaging revealing a dilated, enlarged heart and evidence of cardiac failure (Figure S1A). A left ventricular assist device (LVAD) was inserted for seven weeks, facilitating cardiac recovery and the patient was given vitamin supplementation of riboflavin, thiamine, ascorbic acid, and ubiquinone during this time. At 5 years old, another acute cardiac event triggered readmission with a second LVAD inserted for two weeks and TPN was administered. Cardiac function recovered and was stable for two years. At this point, the patient presented with lethargy and cardiorespiratory insufficiency and a third LVAD was inserted until a donor heart was transplanted a week later, age 7. Clinical follow-up showed no rejection and good cardiac function and the stability of the transplant is maintained on tacrolimus 1mg daily and mycophenolate mofetil 500mg twice daily. Additional clinical complications included developmental delay, persistent left hemiplegia, complete areflexia, and recurrent seizures. Metabolic screening at 7 years demonstrated increased excretion of 3-hydroxyisovaleric acid (3HIA) in urine, and increased 3-hydroxyisovaleryl-carnitine (C5OH) in both urine and a blood spot, consistent with 3-methylcrotonyl-CoA carboxylase and biotinidase deficiency.^(24, 25)^ At 14 years, based on the molecular findings (described below), supplementation was commenced with daily doses of pantothenic acid (550mg), lipoic acid (400mg) and biotin (10mg). Subsequently daily ferrous sulphate (210mg) was added in 2023 after diagnosis of iron deficiency associated with recurrent episodes of loose stools and hair loss. Epilepsy is controlled with twice daily zonesamide (200mg), lamotrigine (225mg), clonazepam (10mg) and brivaracetam (50mg). Infection is suppressed with co-trimoxazole 480mg daily. Assessing the impact of the vitamin supplements has been challenging in an adolescent with multiple medical problems. The parents are excellent observers and consider growth, development and epilepsy control have all improved after six years on supplements. Puberty commenced normally. She continues to have occasional seizures at the time of her periods.

**Figure 1.**
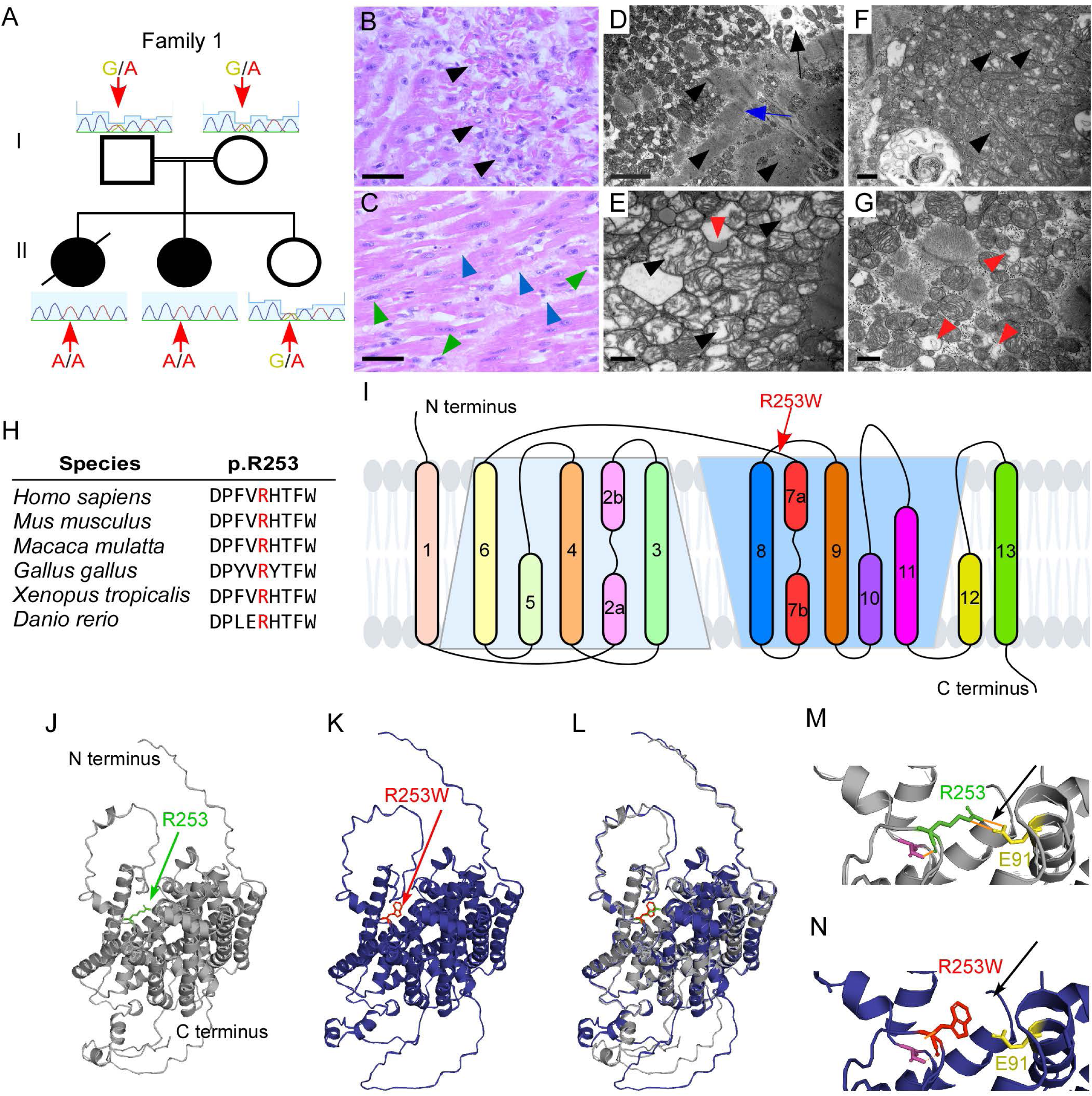
Clinical presentation and modelling of *SLC5A6^R253W^* mutation. (A) Pedigree of Family 1. The unaffected consanguineous parents and child II-3 were heterozygous for the *SLC5A6* mutation (G/A), whereas children II-1 and II-2 both had DCM and were homozygous (A/A). (B-G) Imaging of a left ventricular heart biopsy from child II-2. (B,C) Myocardial disarray with hypertrophied cardiomyocytes (black arrowheads), hyperchromatic nuclei (green arrowheads) and interstitial fibrosis (blue arrowheads). (D-G) Sarcomeric wasting (black arrow in D), sarcomeric disarray (blue arrow in D), Z band disarray (black arrowheads in D), fragmented mitochondria (black arrowheads in E,F) and mitochondrial degradation (red arrowheads in E,G). (H) Amino acid residue R253 is highly conserved across species. (I) Schematic of SMVT. The patient mutation, R253W, is in the hinge region of the transporter. (J-N) Protein modelling of SMVT with R253 indicated (J). When changed to R253W (K), a salt bridge (orange) is removed (black arrow in M,N). Scale bars B,C = 20µm, D = 2µm, E-G = 500nm.

A myocardial biopsy obtained during LVAD insertion showed myocardial disarray, featuring hypertrophied cardiomyocytes (Figure 1B) with large, irregular, hyperchromatic nuclei. There was also evidence of mild interstitial, perimyocyte and subendocardial fibrosis (Figure 1C). Transmission electron microscopy (TEM) images of the left ventricle revealed sarcomeric wasting and disarray. In some areas, the sarcomeres had fragmented, creating white patches, and the organised parallel alignment of the sarcomeres was lost. Z band disarray was evident, indicated by variations in the thickness of the lines. Additionally, large clusters of mitochondria were observed, containing some fragmented mitochondria (Figure 1D-G). The examining pathologist diagnosed cardiomyopathy with no morphological evidence as to the cause.

In Family 2,^(18)^ consanguineous parents from Tunisia, had five children (Figure S1B). Child II-2 died of unexplained multisystemic decompensation and severe cardiac dysfunction at 8 months (genetic testing was not available at that time) and patient II-5 presented with severe cardiac decompensation at 5 months, with a left ventricular EF of 32% (normal >60%; classed as Day 0 on Figure S1C). Treatment with cardiac drugs (adrenaline, milrinone and furosemide) for 3 days resulted in EF improvement to 62% by Day 10. However, by Day 22, the patient had relapsed with an EF of 17%, necessitating intensive cardiac support with an increasing number of cardiac drugs (digoxin, furosemide, spironolactone, carvedilol and lisinopril). Additionally, following an abnormal plasma acylcarnitine’s profile and a persistent increase in 3-hydroxyisovaleric acid in the urine (3-HIA), biotin supplementation (10 mg on Day 22, increased to 15 mg on Day 33) and then pantothenic acid (100 mg from Day 64) were initiated. Following the increase in biotin dose to 15mg, the EF increased to 45% and the patient was maintained on vitamins and four cardiac drugs (digoxin, furosemide, carvedilol and lisinopril) for a further 79 days. The number of cardiac drugs were slowly reduced as the EF rose to a healthy 66% on Day 121 and 75% on Day 150 (Figure S1B). Bi-vitamin supplementation remains ongoing, and the EF has remained within the normal range. The patient also presented with severe hypogammaglobulinemia that interestingly corrected itself after vitamin supplementation.

### Identification of *SLC5A6* mutation and protein modelling

Whole-exome sequencing of Family 1 revealed a novel homozygous missense variant in *SLC5A6* ((GRCh37) Chr2:27427777 G>A) in patients II-1 and II-2. The mutation results in a substitution of arginine at position 253 to a tryptophan (p.R253W). All unaffected family members were heterozygous for the variant (Figure 1A).

The mutation, in exon 9, affects a highly conserved amino acid (Figure 1H), with an allele frequency of 0.00001983 (gnomADv4.1) and a high pathogenicity prediction (CADD score: 27.4). Within the SMVT transporter the R253W mutation is located on the extracellular loop between transmembrane domains 6 and 7, spanning the hinge region of the LeuT fold (Figure 1I). *In silico* protein modelling indicated that the substitution of arginine to tryptophan introduces a large bulky aromatic ring, which is predicted to induce steric hindrance (Figure 1J-L) and prevent the formation of a salt bridge with E91 (Figure 1M,N).

Protein modelling of the P437L mutation (C>T) in *SLC5A6*, again in a highly conserved residue, (Figure S1D,E) reported in Family 2,^(18)^ showed the overall positioning of the large intracellular loop changes and a novel polar bond is formed between L437 and V434 (Figure S1F-J).

Both pathogenic mutations in *SLC5A6*, therefore, predicted to induce conformational changes in the tertiary structure of the protein, and the likely outcome of these mutations is affected vitamin uptake.

### *Slc5a6^cKO^* mice develop cardiomyopathy

Mice constitutively deficient for *Slc5a*6 (Figure S2A) exhibited embryonic lethality, with underdeveloped embryos at embryonic day (E)9.5 and E10.5 (Figure S2B-E). The number of *Slc5a6^Tm1a/Tm1a^* mutant embryos was significantly underrepresented when collected between E9.5 and E15.5 (Figure S2F).

SLC5A6 was confirmed to be strongly expressed in the mouse and human embryonic heart (Figure S2H-K). A cardiac-specific *Slc5a6* knockout (KO) (*Slc5a6^cKO^*) mouse line was produced by crossing conditional *Slc5a6^Tm1c^* mice with the *Tnt-Cre* line (Figure S2A). RT-PCR confirmed heart-specific deletion of exons 7-10 (Figure S2L) resulting in a premature stop codon (Figure S2A). The *Slc5a6^cKO^* mice were observed at the expected numbers at weaning (Figure S2M).

Histological analysis at E15.5 revealed that *Slc5a6^cKO^*embryos displayed no developmental cardiac abnormalities (Figure 2A,B). *Slc5a6^cKO^*mice were phenotypically normal at birth and into early adulthood, but by 20 weeks there was a significant decrease in body weight (*p*=0.0195; Figure S2N). Sudden death occurred in *Slc5a6^cKO^*mice at 26 weeks (*n*=3), prompting euthanasia of subsequent litters by 20 weeks. Histological analysis of the hearts revealed severe cardiac dilation, with enlarged atria and ventricles (Figure 2C-F). At 14 weeks cardiomyocyte hypertrophy was observed in both the right and left ventricles (LV *p*=0.0016, RV *p*=0.0067; Figure 2G-I). No differences were observed in heart weight to body weight or to tibia length ratios (Figures 2J; S2O), suggesting the heart is dilated with a larger internal volume but no increase in mass. Fibrosis was significantly increased 3-fold throughout the ventricular walls of the *Slc5a6^cKO^* hearts compared to control hearts (*p*=0.0007; Figure 2K-M). An increase in atrial natriuretic peptide (*Nppa*; *p*=0.037) and a decrease in alpha-cardiac myosin heavy chain (*Myh6*; *p*=0.0432) expression were observed in *Slc5a6^cKO^*hearts at 20 weeks (Figure S2P,Q).

**Figure 2.**
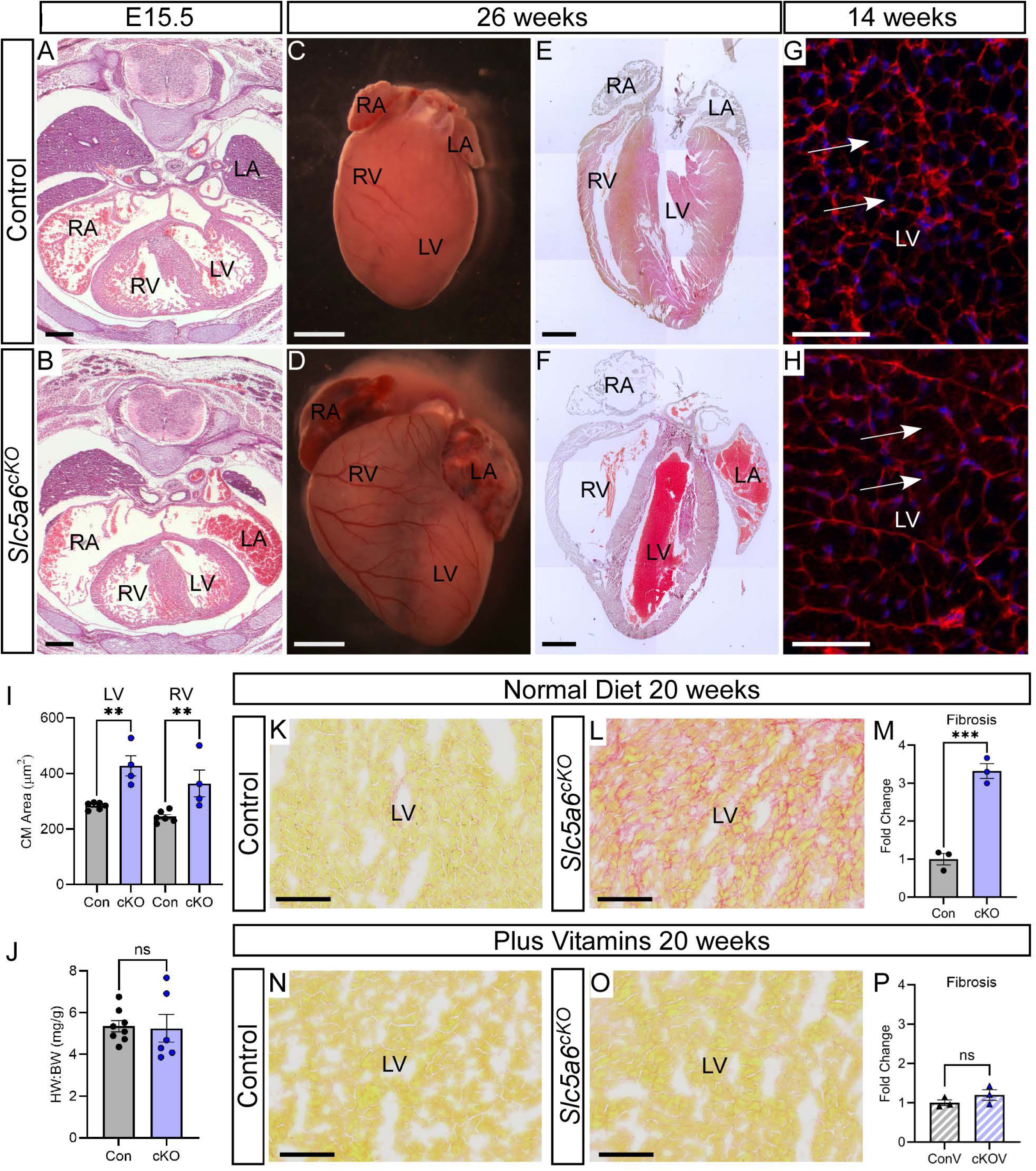
Cardiac-specific *Slc5a6* KO (*Slc5a6^cKO^*) mouse recapitulates DCM phenotype. (A,B) Hearts from E15.5 control embryos (A) were comparable with *Slc5a6^cKO^* mutant embryos (B). (C-F) *Slc5a6^cKO^* mutant hearts collected at 26 weeks were visibly larger and dilated (D) with ventricular and atrial enlargement (F) compared to a controls (C,E). (G,H) WGA staining was performed on coronal heart sections of 14-week control (G) and *Slc5a6^cKO^*mutant hearts (H). (I) Cardiomyocytes were significantly larger in both ventricles in the *Slc5a6^cKO^*mutant (*n*=4) compared to the control (*n*=6) hearts (cell area measured from four regions of interest per ventricle with three technical repeats per heart). (J) No difference in heart weight (HW) to body weight (BW) ratio was observed between control (*n*=8) and *Slc5a6^cKO^* mutant (*n*=6) mice at 20 weeks. (K-P) Fibrosis was increased in *Slc5a6^cKO^* mutant hearts (*n*=3; L,M) compared to control hearts (*n*=3; K,M) at 20 weeks. (N-P) Following vitamin supplementation, no increase in fibrosis in *Slc5a6^cKO^*mutants (*n*=3) compared to control (*n*=3) mice was observed. Data are represented as mean ± SEM. ns = nonsignificant, **p < 0.01, ***p < 0.001 by one-way ANOVA with Bonferroni correction for multiple comparisons or unpaired t-test for two sample comparisons. RV, right ventricle; LV, left ventricle; RA, right atria; LA, left atria. Con, control; cKO, *Slc5a6^cKO^*; ConV, vitamin-supplemented control; cKOV, vitamin-supplemented *Slc5a6^cKO^*. Scale bars A,B = 500µm, C,D = 2mm, E,F = 1mm, G,H = 50µm, K,L,N,O = 100µm.

Thus, the *Slc5a6^cKO^*mice develop the key features of DCM with a dilated heart, cardiomyocyte hypertrophy, fibrosis and fetal gene reprogramming.

### Cardiac function is severely reduced in *Slc5a6^cKO^* mice

Cardiovascular Magnetic Resonance (CMR) imaging and three-lead ECG readings were used to assess the impact of cardiac structural abnormalities on cardiac function. CMR performed at 20 weeks demonstrated significantly impaired cardiac function in the *Slc5a6^cKO^* mutants (*n*=4), with significant decreases in EF (*p*=0.0207; 28.8% ±12.29 compared to 61% ±19.72 in controls (*n*=6)), stroke volume (SV, *p*=0.0146; 16.1µl ±6.74 compared to 34.4µl ±10.3 in controls), and cardiac output (CO, *p*=0.007; 6.8ml/min ±2.1 compared to 15.8ml/min ±4.63 in controls) (Figure 3A-C).

**Figure 3.**
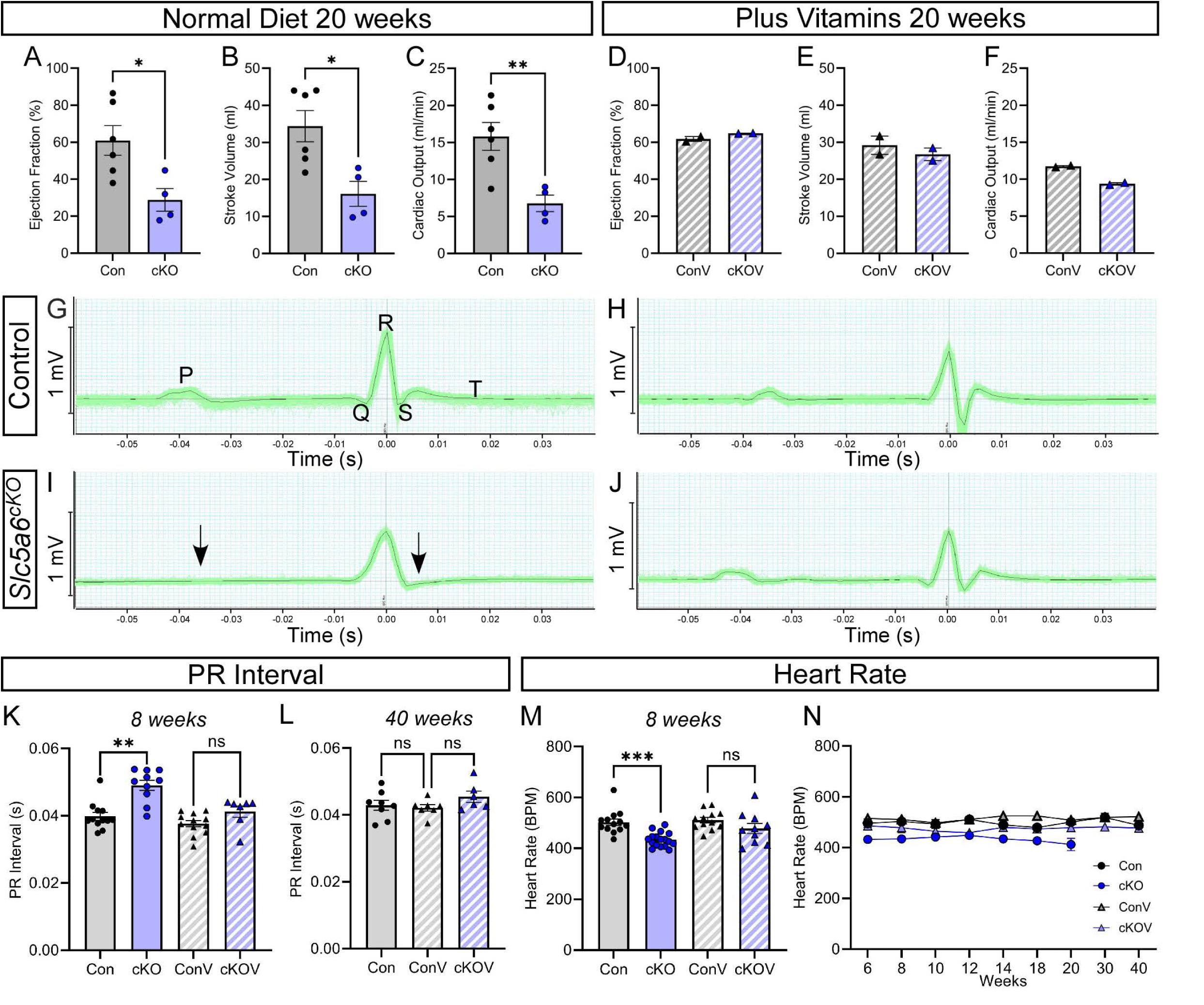
Vitamin supplementation prevents cardiac function defects in *Slc5a6^cKO^* mutant mice. (A-F) CMR imaging at 20 weeks. A significant decrease in ejection fraction (A), stroke volume (B) and cardiac output (C) was observed in *Slc5a6^cKO^*mutants (*n*=4) compared to controls (*n*=6). (D-F) After vitamin supplementation, no apparent differences in cardiac function was seen (*n*=2 for each genotype). (G-J) Representative ECG traces from 20-week mice. Control (G) and vitamin-supplemented control (H) mice have well-defined P, QRS and ST parameters whereas *Slc5a6^cKO^*mice (I) present with an undefined and relatively flat trace, with loss or reduction of the P wave and an undefined ST wave (arrows). ECG traces from vitamin-supplemented *Slc5a6^cKO^* mutants (J) were comparable to control mice. (K,L) The PR interval was measured and was significantly longer in *Slc5a6^cKO^* compared to control mice. No difference was observed in vitamin-supplemented *Slc5a6^cKO^*mice at 8 and 40 weeks (K,L; 8 weeks: Con *n*=12, cKO *n*=10, ConV *n*=12, cKOV *n*=6; 40 weeks: Con *n*=8, ConV *n*=7, cKOV *n*=6). (M) The heart rate was significantly lower in *Slc5a6^cKO^* mice at 8 weeks (*n*=17) compared to controls (*n*=13) and no differences were seen in vitamin-supplemented mice (ConV *n*=12 and cKOV *n*=10). (N) Longitudinal measurements illustrated the heart rate of *Slc5a6^cKO^*mice was consistently lower than control and vitamin-supplemented *Slc5a6^cKO^*mice. Con, control; cKO, *Slc5a6^cKO^*; ConV, vitamin-supplemented control; cKOV, vitamin-supplemented *Slc5a6^cKO^*. Data are represented as mean ± SEM. ns = nonsignificant, **p < 0.01, ***p < 0.001 by one-way ANOVA with Bonferroni correction for multiple comparisons or unpaired t-test for two sample comparisons.

ECG was performed from 8 to 20 weeks on control and *Slc5a6^cKO^*mice to monitor cardiac function in vivo. However, due to the natural variation between mouse heart rhythms and progression of disease with time, not all parameters could be accurately quantified at each time point from the average ECG traces. For example, at 20 weeks there was a clear visual difference between the ECG traces from control and *Slc5a6^cKO^* mice, with the flattened *Slc5a6^cKO^* trace with no clearly defined ST wave (Figure 3G,I). Data was, therefore, tabulated to show the percentage of mice for which each parameter could be calculated (Table 1). This demonstrated an overall defect in cardiac conduction from 8 weeks which deteriorated severely by 20 weeks, which was further emphasized as ECG traces were only quantifiable in 7.1% of *Slc5a6^cKO^* mice compared to 100% of controls.

**Table 1.**
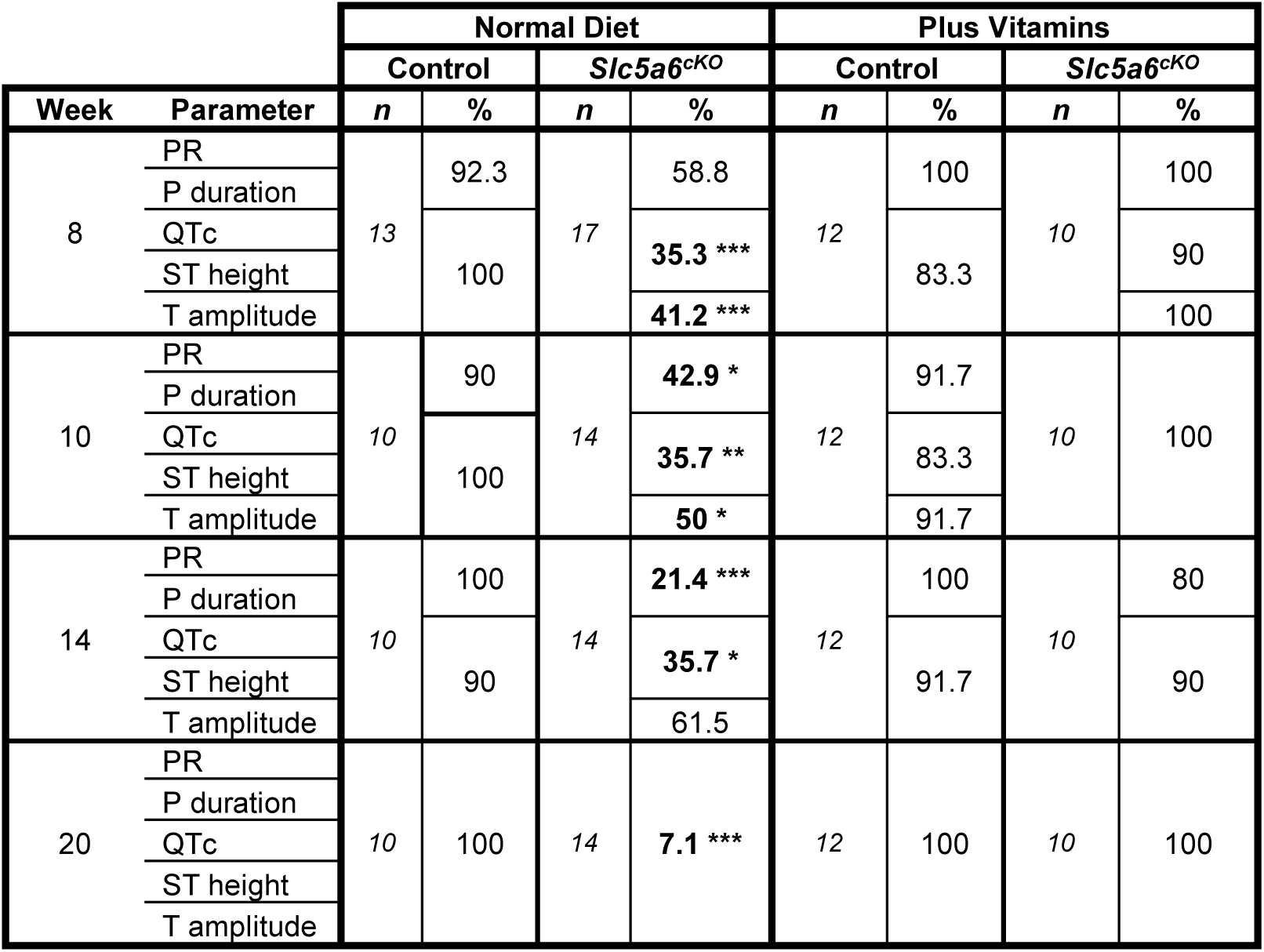
Summary of quantifiable ECG parameters at different time points. ECG traces were averaged for the third minute, and the following parameters were calculated for each mouse: PR interval, P duration, QTc interval, ST height and T amplitude. As some parameters could not be quantified the association between the number of quantifiable and non-quantifiable parameters at each time point for each genotype was calculated. *p<0.05, **p < 0.01, ***p < 0.001 (Fishers Exact test).

Analysis of the quantifiable ECG parameters at 8 weeks showed a significant increase in PR interval (*p*=0.004; Figure 3K, control *n*=12, *Slc5a6^cKO^ n*=10) and a decrease in T amplitude (*p*=0.0483; Figure S3B, control *n*=13, *Slc5a6^cKO^ n*=7) in *Slc5a6^cKO^*mice. Heart rate was consistently lower in the *Slc5a6^cKO^* mice (*p*=0.0004; Figure 3M, control *n*=13, *Slc5a6^cKO^ n*=17).

The cardiac-specific deletion of *Slc5a6*, therefore, caused anatomical, histological and functional cardiac alterations typical of DCM, resulting in sudden death by 26 weeks. Thus, the *Slc5a6^cKO^* mouse is a good model of DCM, recapitulating the DCM seen in the patients.

### *Slc5a6^cKO^* mice have reduced vitamin transport

To establish the cause of DCM, mass spectrometry metabolomic analyses of 8-week *Slc5a6^cKO^*mice was performed and compared with control mice. Biotin, a cofactor for carboxylases, and pantothenic acid, required for the synthesis of CoA, are both transported by SMVT.^(26, 27)^ FIA-MS/MS analysis of plasma acylcarnitine revealed a significant increase in 3-hydroxyisovaleryl-carnitine (C5OH) (*p*=0.0002; Figure 4A), a hallmark of 3-methylcrotonyl-CoA carboxylase (MCC) and biotinidase deficiency. LC-MS-qTOF analysis on heart powder detected significantly increased levels of 3-hydroxyisovaleric acid (3-HIA; *p*<0.0001) and 2-methylcitric acid (2MCA; *p*=0.0017), indicative of MCC and propionyl-CoA carboxylase (PCC) deficiencies, respectively (Figure 4B,C). Additionally, cardiac levels of pantothenic acid (*p*<0.0001), and its derivatives phosphopantothenic acid (*p*<0.0001) and phosphopantethein (*p*=0.0219) were significantly decreased, leading to impaired CoA synthesis, as evidenced by reduced CoA-glutathione (*p*=0.0002) (Figure 4D-G). Western blot analysis confirmed a significant reduction in MCC and PCC biotinylation in *Slc5a6^cKO^* hearts at 5 weeks (before any structural or functional cardiac differences were observed; *p*=0.0178) and 20 weeks (experimental endpoint; *p*=0.0006) (Figures 4H,J; S4A,C), which was unchanged in the liver (Figures 4I,K; S4B,D). Pyruvate carboxylase (PC) levels were unaffected (Figures 4M-P; S4A-D). Thus, these results indicated biochemically there was a heart-specific reduction in biotin and pantothenic acid transport in *Slc5a6^cKO^*mice.

**Figure 4.**
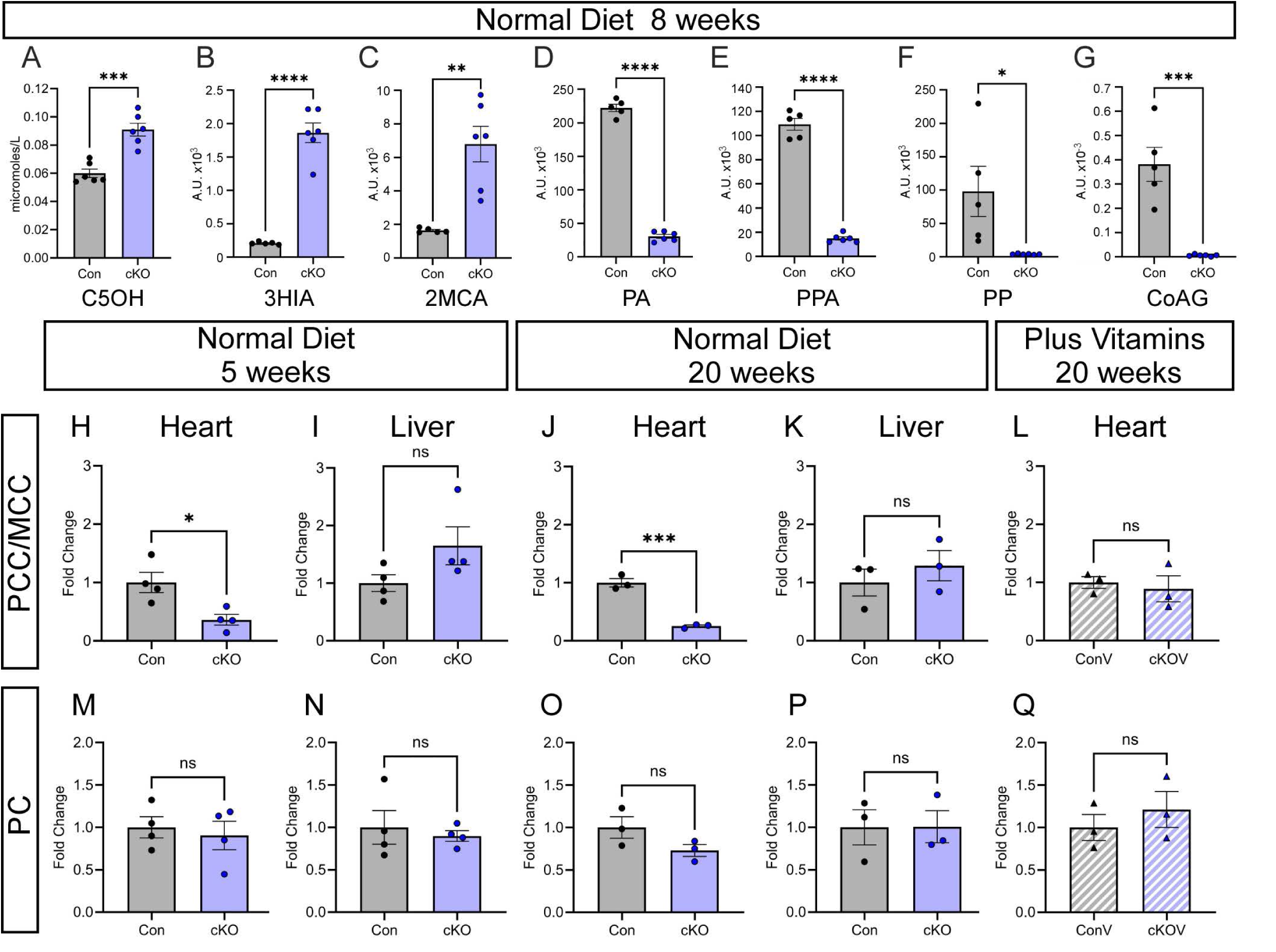
Reduced vitamin transport in *Slc5a6^cKO^* hearts. (A-G) Acylcarnitine analysis of plasma showed a significant increase in C5OH in *Slc5a6^cKO^* mutants (*n*=6) compared to controls (*n*=6). Metabolic analysis of heart powder showed significant changes in 3HIA, 2MCA, PA, PPA, PP and CoAG in *Slc5a6^cKO^* mutants (*n*=5) compared to controls (*n*=6). (H-Q) A decrease in PCC/MCC biotinylation was observed in *Slc5a6^cKO^* mutants at 5 and 20 weeks (H,J), with no corresponding difference in liver samples (I,K). In vitamin-supplemented *Slc5a6^cKO^* mutants PCC/MCC biotinylation levels were comparable to control levels (L). No change in PC was observed in the hearts or livers (M-Q). Data are represented as mean ± SEM. ns, nonsignificant, *p < 0.05, **p < 0.01, ***p < 0.001, ****p < 0.0001 by one-way ANOVA with Bonferroni correction for multiple comparisons or unpaired t-test for two sample comparisons. C5OH, 3-hydroxyisovaleryl-carnitine; 3-HIA, 3-hydroxyisovaleric acid; 2MCA, 2-methylcitric acid; PA, pantothenic acid; PPA, phosphopantothenic acid; PP, phosphopantethein; CoAG, CoA-glutathione; Con, control; cKO, *Slc5a6^cKO^*; ConV, vitamin-supplemented control; cKOV, vitamin-supplemented *Slc5a6^cKO^*.

### Vitamin supplementation ameliorates the DCM phenotype

As patient II-5 in Family 2 responded well to early vitamin treatment, to specifically assess the impact of vitamin supplementation on heart function, biotin and pantothenic acid were added to the water and diet of *Slc5a6^cKO^* mice at doses equivalent to those used in humans (Supplementary Table 1).^(28)^ Supplementation was initiated in parental mice before conception, continued throughout gestation and provided long-term to the offspring. Maintenance of normal levels of MCC and PCC biotinylation in 20-week hearts of supplemented *Slc5a6^cKO^* mice (Figure 4L) indicated successful vitamin uptake.

Vitamin-supplemented *Slc5a6^cKO^* mice survived to 40 weeks, at which time the mice were electively culled. Vitamin supplementation did not affect body weight at 20 or 40 weeks (Figure S5A-F) and it prevented the excessive deposition of fibrosis in *Slc5a6^cKO^* hearts (Figures 2N-P; S5G-K) and there was no longer altered expression of *Nppa* and *Myh6* at 20 weeks (Figure S2R,S). There were no discernible differences in cardiac function; EF, SV or CO between vitamin-supplemented control and *Slc5a6^cKO^* mice (Figure 3D-F), and all ECG traces could be quantified at 20 weeks (Figure 3H,J; Table 1). No significant difference in any ECG parameter was observed at any time point (Figures 3K-N; S3A-E). Vitamin supplementation, therefore, prevented all the hallmarks of DCM observed in *Slc5a6^cKO^* mice.

### Mitochondrial complexity increases with disease progression in *Slc5a6^cKO^* mice

As changes in mitochondria morphology are associated with metabolic dysfunction and disease progression,^(29, 30)^ we analysed the ultrastructure of cardiac mitochondria using TEM.

At 20 weeks, control hearts presented with predominantly spherical shaped mitochondria, with minimal branching or complex shapes, and formed parallel clusters along longitudinal myofibers (Figures 5A,C; S7A-D). In contrast, mitochondria in *Slc5a6^cKO^* hearts were irregularly shaped and arranged randomly between myofibrils with evidence of mitochondrial degradation (Figures 5B,D; S7E-L). The mitochondrial complexity index (MCI) was significantly increased in *Slc5a6^cKO^* hearts (*p*<0.0001) with a corresponding significant decrease in volume (*p*<0.0001; Figure 5I,J; Table 2). The *Slc5a6^cKO^* hearts, therefore, had smaller mitochondria, with more complex shapes including excessive elongation, branching and nanotunnels (Figure S7M-P). This is further represented by the significant difference in the lines of best fit for controls versus *Slc5a6^cKO^*hearts on the bivariate plot (*p*<0.0001; Figure 5K) and a shift to the right in the cumulative frequency distributions (Figure 5L).

**Figure 5.**
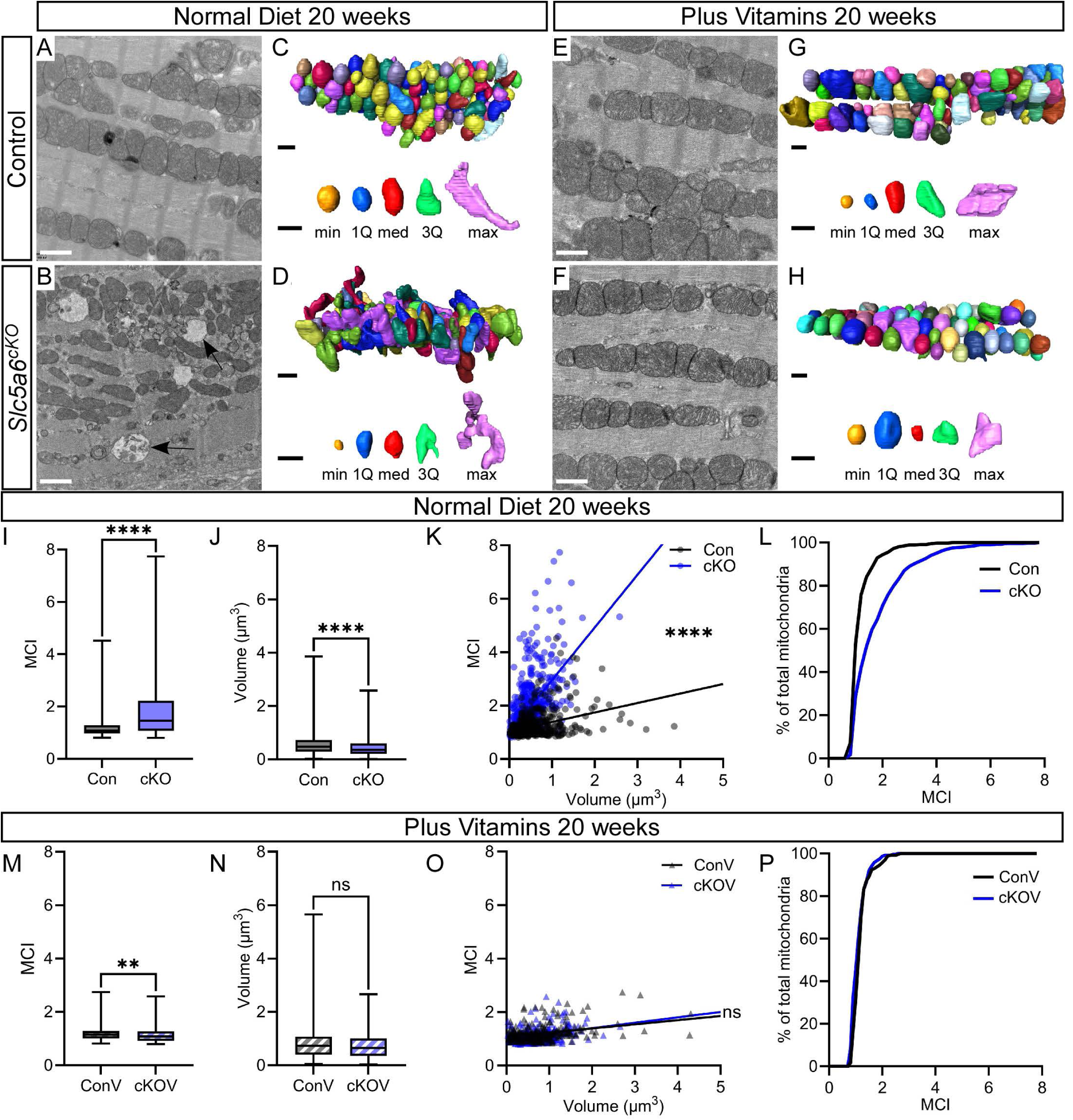
Increased mitochondria complexity following loss of *Slc5a6* is prevented by vitamin supplementation. (A-H) Mitochondria imaging at 20 weeks by TEM (A,B,E,F) and 3D reconstructions (C,D,G,H). Mitochondria in controls, vitamin-supplemented controls and vitamin-supplemented *Slc5a6^cKO^* hearts were all comparable (A,C,E-H) whereas mitochondria in *Slc5a6^cKO^* hearts (B,D) were unevenly arranged and irregularly shaped with evidence of mitochondrial degradation (arrows in B). Representative individual mitochondria of the minimum (min), first quartile (1Q), median (med), third quartile (3Q) and maximum (max) MCI are shown (total number of mitochondria analysed: Con = 468, cKO = 530, ConV = 241 and cKOV = 240). (I-L) The MCI was significantly increased in 20-week *Slc5a6^cKO^* mice on a normal diet (I) and the mitochondrial volume was significantly decreased (J). Bivariate analysis of MCI:volume shows a significant difference in the regression line (K) and the cumulative frequency distribution curves do not overlap (L). (M-P) In vitamin-supplemented *Slc5a6^cKO^* mice the MCI was significantly reduced (M) and there was no difference in mitochondrial volume (N) and complexity (O,P). Data are represented as mean ± SEM. ns, nonsignificant, ***p < 0.01, ****p < 0.0001, by unpaired t-test for two sample comparisons. Con, control; cKO, *Slc5a6^cKO^*; ConV, vitamin-supplemented control; cKOV, vitamin-supplemented *Slc5a6^cKO^*. Scale bars A,B,E,F = 500nm, C,D,G,H = 1µm.

**Table 2.**
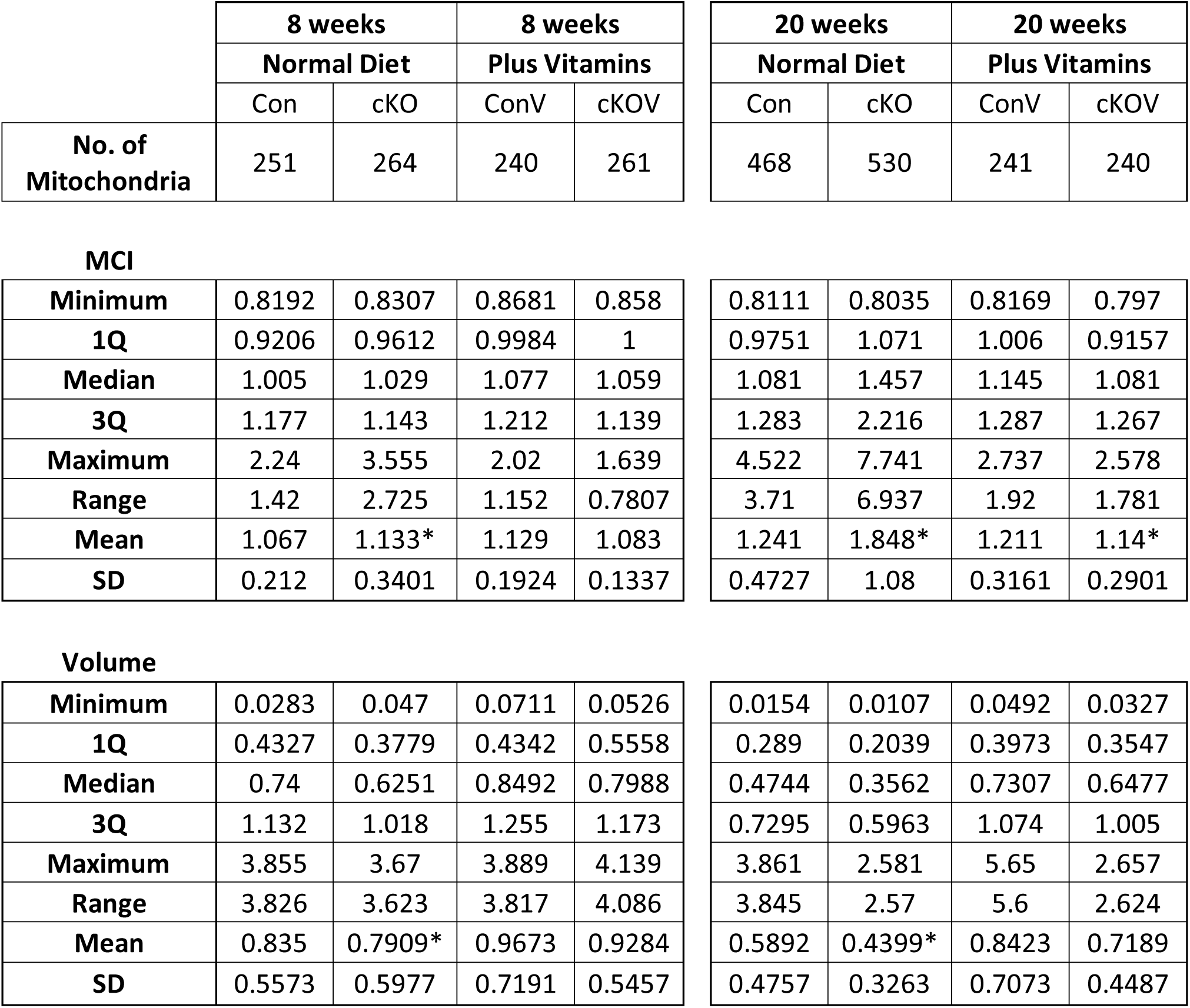
Measurements from mitochondrial reconstructions. Cardiac mitochondrial MCI and volume measurements from normal diet and vitamin-supplemented control and *Slc5a6^cKO^*mice at 8 and 20 weeks. * Indicates the significant differences shown in Figure 5 and Figure S6.

|  | 8 weeks |  | 8 weeks |  | 20 weeks |  | 20 weeks |  |
| --- | --- | --- | --- | --- | --- | --- | --- | --- |
|  | Normal Diet |  | Plus Vitamins |  | Normal Diet |  | Plus Vitamins |  |
|  | Con | cKO | ConV | cKOV | Con | cKO | ConV | cKOV |
| **No. of Mitochondria** | 251 | 264 | 240 | 261 | 468 | 530 | 241 | 240 |
| **MCI** |  |  |  |  |  |  |  |  |
| **Minimum** | 0.8192 | 0.8307 | 0.8681 | 0.858 | 0.8111 | 0.8035 | 0.8169 | 0.797 |
| **1Q** | 0.9206 | 0.9612 | 0.9984 | 1 | 0.9751 | 1.071 | 1.006 | 0.9157 |
| **Median** | 1.005 | 1.029 | 1.077 | 1.059 | 1.081 | 1.457 | 1.145 | 1.081 |
| **3Q** | 1.177 | 1.143 | 1.212 | 1.139 | 1.283 | 2.216 | 1.287 | 1.267 |
| **Maximum** | 2.24 | 3.555 | 2.02 | 1.639 | 4.522 | 7.741 | 2.737 | 2.578 |
| **Range** | 1.42 | 2.725 | 1.152 | 0.7807 | 3.71 | 6.937 | 1.92 | 1.781 |
| **Mean** | 1.067 | 1.133* | 1.129 | 1.083 | 1.241 | 1.848* | 1.211 | 1.14* |
| **SD** | 0.212 | 0.3401 | 0.1924 | 0.1337 | 0.4727 | 1.08 | 0.3161 | 0.2901 |
| **Volume** |  |  |  |  |  |  |  |  |
| **Minimum** | 0.0283 | 0.047 | 0.0711 | 0.0526 | 0.0154 | 0.0107 | 0.0492 | 0.0327 |
| **1Q** | 0.4327 | 0.3779 | 0.4342 | 0.5558 | 0.289 | 0.2039 | 0.3973 | 0.3547 |
| **Median** | 0.74 | 0.6251 | 0.8492 | 0.7988 | 0.4744 | 0.3562 | 0.7307 | 0.6477 |
| **3Q** | 1.132 | 1.018 | 1.255 | 1.173 | 0.7295 | 0.5963 | 1.074 | 1.005 |
| **Maximum** | 3.855 | 3.67 | 3.889 | 4.139 | 3.861 | 2.581 | 5.65 | 2.657 |
| **Range** | 3.826 | 3.623 | 3.817 | 4.086 | 3.845 | 2.57 | 5.6 | 2.624 |
| **Mean** | 0.835 | 0.7909* | 0.9673 | 0.9284 | 0.5892 | 0.4399* | 0.8423 | 0.7189 |
| **SD** | 0.5573 | 0.5977 | 0.7191 | 0.5457 | 0.4757 | 0.3263 | 0.7073 | 0.4487 |

To establish if the changes in mitochondria morphology were associated with early indicators of disease, the mitochondria were investigated at 8 weeks. The overall arrangement of the parallel clusters of the mitochondria along the myofibrils was similar between control and *Slc5a6^cKO^* mice (Figure S6A,B). There was, however, a significant increase in the MCI of mitochondria in *Slc5a6^cKO^* hearts compared to controls (*p*=0.0075), although there was no change in volume (Figure S6C,D,I-L; Table 2).

Cardiac mitochondria morphology was then assessed in vitamin-supplemented mice. At 20 weeks the morphology of mitochondria was similar between control and *Slc5a6^cKO^* hearts, and there was a significant reduction in MCI (*p*=0.0013) but no change in volume (Figures 5E-H,M,N; Table 2), suggesting that vitamins prevented the abnormal mitochondrial morphology. The bivariate plot of mitochondrial MCI:volume and cumulative MCI frequency showed no differences between vitamin-supplemented control and *Slc5a6^cKO^*hearts at 20 weeks (Figure 5O,P). At 8 weeks there was a small yet significant decrease in MCI (*p*=0.0462) but not volume (Figure S6E-H,M,N; Table 2). Interestingly, in vitamin-supplemented mice at 8 weeks, the correlation between the spread of the mitochondria was similar but still significantly different (*p*=0.0084; Figure S6O,P).

The subtle change in mitochondria complexity at 8 weeks coincides with the early indication of cardiac dysfunction in *Slc5a6^cKO^* mice. As DCM progresses towards heart failure at 20 weeks, the mitochondria become more complex in shape and is associated with a corresponding increase in mitochondria degradation. Vitamin-supplementation mitigates this effect in *Slc5a6^cKO^* mice.

### Altered metabolism pathways underlie the onset of DCM in *Slc5a6^cKO^* mice

To elucidate the cellular mechanism driving DCM progression in the *Slc5a6^cKO^* mice, we performed an unbiased proteomics analysis to identify altered signalling pathways at 8 weeks before the overt DCM phenotype was evident. Protein was extracted from control and *Slc5a6^cKO^* hearts, from normal diet and vitamin-supplemented animals (*n*=5 per group).

Analysis of differential protein expression, using principal component analysis and a hierarchical heat map plot, showed that the *Slc5a6^cKO^* samples formed a distinct group compared to the control and vitamin-supplemented samples (Figure 6A,B).

**Figure 6.**
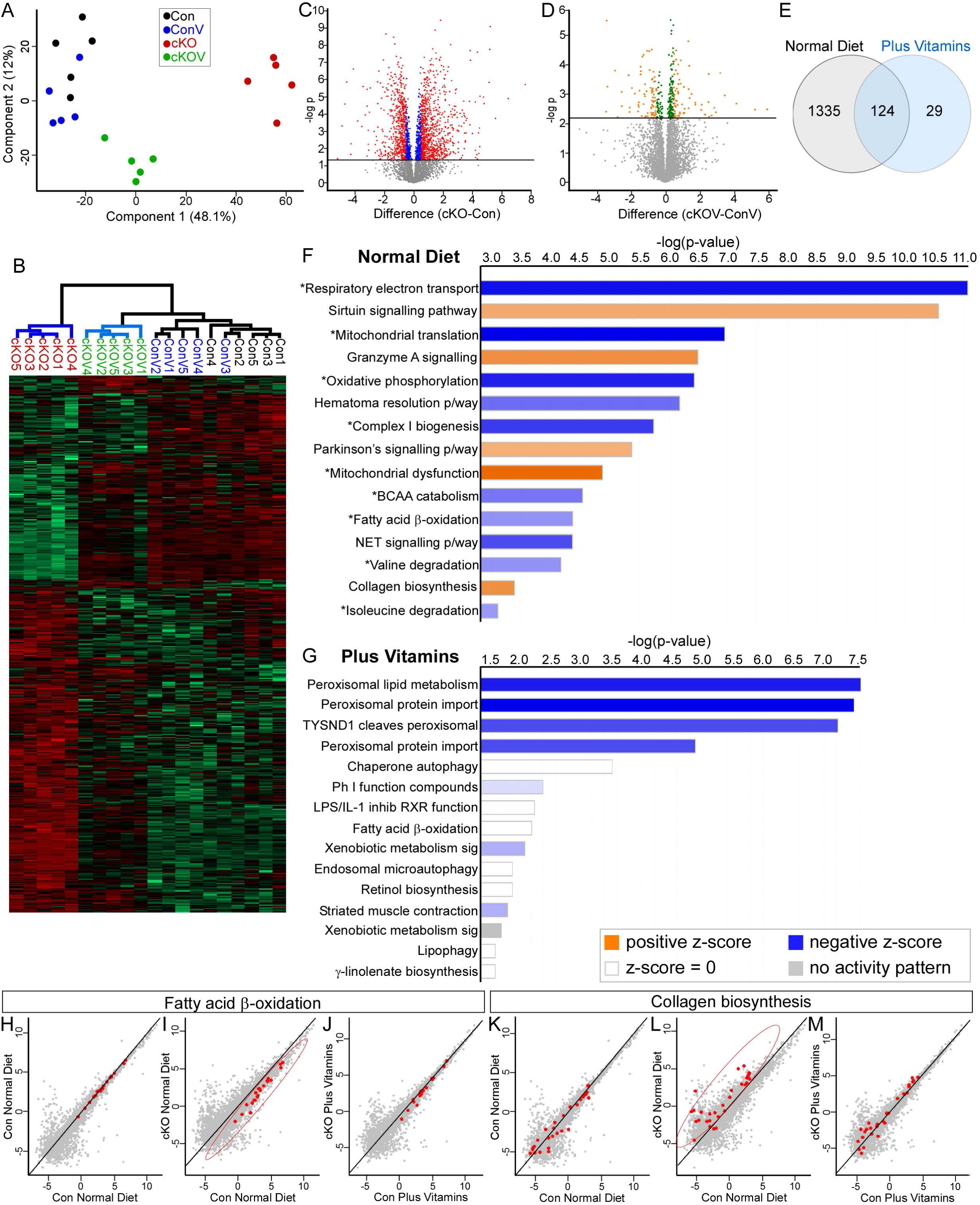
Proteomic analysis revealed altered metabolism and increased fibrosis correlate with early indicators of DCM in *Slc5a6^cKO^* mice. (A-E) Proteomic analysis at 8 weeks. PCA plot (A) and heatmap (B) illustrate that the proteome of the *Slc5a6^cKO^* mutants was distinct from the three other groups (control, vitamin-supplemented control and vitamin-supplemented *Slc5a6^cKO^* hearts; *n*=5 per group). (C,D) Volcano plots showing proteins with a significant (q<0.05) fold change. (C) Under a normal diet, 1459 proteins with a fold change of ≤-0.5 and ≥0.5 (red dots) and 478 proteins with a fold change of >-0.5 and <0.5 (blue dots). (D) With vitamin supplementation only 153 proteins with a fold change of ≤-0.5 and ≥0.5 (orange dots) and 179 proteins with a fold change of >-0.5 and <0.5 (green dots) were seen. (E) Venn diagram illustrating 124 of the proteins with a fold change of ≤-0.5 and ≥0.5, remained significantly differentially expressed following vitamin supplementation. (F,G) IPA analysis of up- and down-regulated proteins. For the mice on a normal diet, the majority of the enriched down-regulated pathways in *Slc5a6^cKO^* mutant mice are linked to different components of the energy metabolism pathways in mitochondria (asterisks) (F). In comparison, vitamin-supplemented *Slc5a6^cKO^* mutant mice only had downregulation of peroxisome-related pathways (G). (H-M) Correlation graphs of proteins involved in fatty acid β-oxidation (H-J) or collagen biosynthesis (K-M) (red dots) illustrate the shift away from the line of best fit in the *Slc5a6^cKO^* mutants (circled dots), which shift back to the line following vitamin supplementation.

For normal diet, a total of 1459 proteins showed significant differential expression (599 downregulated, 860 upregulated; *p*<0.05) in *Slc5a6^cKO^* hearts compared to controls (Figure 6C). For the vitamin-supplemented mice, only 153 proteins showed significant differential expression (81 downregulated, 72 upregulated; *p*<0.05) in *Slc5a6^cKO^* hearts compared to controls (Figure 6D). Of the proteins differentially expressed under a normal diet, only 124 of those proteins remained significantly differentially expressed following vitamin supplementation (Figure 6E). There was no significant difference in protein expression between the normal diet and vitamin-supplemented control samples.

Pathway analysis was performed on the significantly down- and up-regulated proteins combined. From mice on a normal diet, in the top 15 significantly enriched pathways (Figure 6F), the upregulated pathways included the ‘sirtuin signalling pathway’ and ‘collagen biosynthesis and modifying enzymes’, which are both linked to increasing fibrosis,^(31, 32)^ and ‘Granzyme A signalling’ which correlates with the upregulation of mitochondrial dysfunction pathways.^(33)^ There was downregulation of key pathways in energy metabolism: ‘respiratory electron transport’, ‘mitochondrial translation’, ‘oxidation phosphorylation’, ‘complex I biogenesis’, ‘branched-chain amino acid catabolism (BCAA)’ (specifically valine and isoleucine degradation), and ‘fatty acid beta-oxidation’. Hematome resolution signalling pathway is linked to genes involved in peroxisomal beta-oxidation.^(34)^ In contrast, the dramatically reduced number of enriched pathways in the vitamin-supplemented hearts (Figure 6G), highlighted there was only downregulation of peroxisome related pathways.

Correlation analyses illustrated that key metabolic pathways were affected following vitamin supplementation. Proteins associated with the fatty acid β-oxidation pathway, which were downregulated, clustered below the line of best fit in *Slc5a6^cKO^* hearts, then aligned with control levels in vitamin-supplemented *Slc5a6^cKO^* hearts (Figures 6H-J). Similarly, collagen biosynthesis proteins, which were upregulated in *Slc5a6^cKO^* hearts, aligned with control levels after supplementation (Figures 6K-M).

Energy metabolism involves several interconnected pathways. Analysis of different pathways showed that the majority of proteins linked to the TCA cycle, fatty acid metabolism, BCAA catabolism and the electron transport chain were all downregulated in the *Slc5a6^cKO^* hearts on a normal diet, with a corresponding increase in proteins involved in glycolysis (Figure 7A), which is a common compensatory response in heart failure.^(35)^

**Figure 7.**
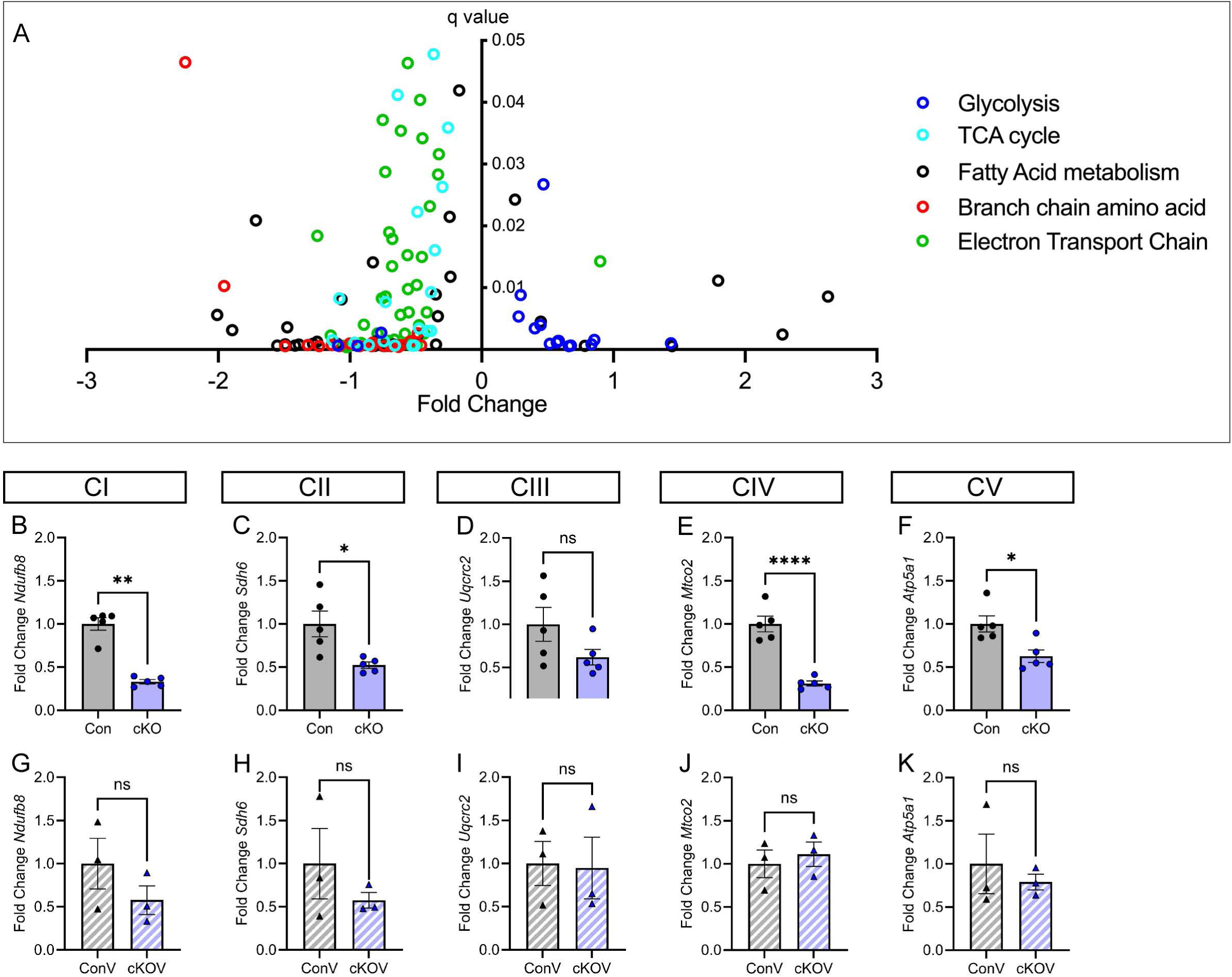
Decreased energy metabolism pathways and the defective electron transport chain in *Slc5a6^cKO^*mice. (A) Summary of five different pathways involved in energy metabolism identified in the proteomics analysis of 8-week hearts. The majority of proteins in the TCA cycle, fatty acid metabolism, branch chain amino acid catabolism and electron transport chain are downregulated in *Slc5a6^cKO^* mutants, with an increase in proteins associated with glycolysis. (B-K) qPCR analysis of genes representing a subunit from each complex of the electron transport chain (CI: *Ndufb8*, CII: *Sdhb*, CIII: *Uqcrc2*, CIV: *Mtco2* and CV: *Atp5a1*) using 8-week hearts. There was a significant downregulation in Complexes I, II, IV, and V in *Slc5a6^cKO^*hearts (*n*=5 hearts for each genotype; B-F) and no corresponding decrease in vitamin-treated hearts (*n*=3 hearts for each genotype; G-K). Data are represented as mean ± SEM. ns, nonsignificant, *p < 0.05, **p < 0.01, ****p < 0.0001 by unpaired t-test. Con, control; cKO, *Slc5a6^cKO^*; ConV, vitamin-supplemented control; cKOV, vitamin-supplemented *Slc5a6^cKO^*.

Each mitochondrial complex (C) in the electron transport chain, CI to CV, is made from multiple subunits. Notably, from the subunits detected in the proteomics experiment, the components exhibited widespread reduced expression in *Slc5a6^cKO^*mutants compared to controls, with the following percentage of subunits showing decreased expression: CI 90.2%, CII 100%, CIII 77.8%, CIV 77%, CV 76.5% (q<0.05, all FC; Supplementary Table 2). Remarkably, vitamin supplementation substantially prevented these deficits, reducing the proportion of downregulated subunits to only two in CI (NDUFA11 and NDUFS6) and one in CIV (MT-CO3) in vitamin-supplemented *Slc5a6^cKO^*hearts compared to controls. These findings were further validated by qPCR analysis of representative subunits of each complex highlighted in Supplementary Table 2, which confirmed significant downregulation in Complexes I (*p*=0.0079), II (*p*=0.0142), IV (*p*<0.0001), and V (*p*=0.0141) in *Slc5a6^cKO^* hearts (Figure 7B-F), with no significant differences detected in vitamin-treated hearts (Figure 7G-K).

Overall, these findings demonstrate that *Slc5a6* deficiency disrupts the cardiac proteome homeostasis, leading to metabolic defects and extracellular matrix remodelling. Importantly, vitamin supplementation effectively mitigates these molecular perturbations, maintaining proteomic profiles at control levels and preventing DCM progression.

## Discussion

In this study, we identified *SLC5A6* mutations as a novel genetic cause of dilated cardiomyopathy (DCM) in two unrelated families. Clinical outcomes in these families highlight the critical importance of early intervention: timely and sustained vitamin supplementation prevented the progression to end-stage heart failure and the need for heart transplantation, whereas delayed treatment was associated with irreversible cardiac decline requiring transplantation. Using a cardiac deletion mouse model of *Slc5a6*, *Slc5a6^cKO^*, we confirmed that *Slc5a6* is a novel DCM causative gene linked to abnormal mitochondria and impaired energy metabolism which are prevented with vitamin supplementation.

The clinical symptoms in patients with *SLC5A6* mutations typically manifest between one week to 15 years, with 18% of patients dying with no reported treatment. It is an autosomal recessive disease, but with no obvious genotype:phenotype correlation. A third of patients exhibit cardiac complications including ventricular fibrillation,^(13)^ left ventricle dysfunction,^(18)^ non-specific ST and T wave abnormalities^(12, 22)^ and DCM, requiring heart transplantation.^(14, 17)^

In *Slc5a6^cKO^* mice, inactivation of *Slc5a6* specifically in cardiomyocytes results in a lethal phenotype with characteristic anatomical, functional and histological features consistent with DCM.^(36–38)^ The *Slc5a6^cKO^* mice were phenotypically normal up to 8 weeks when abnormal ECG readings were identified, demonstrating early indicators of DCM.^(39, 40)^ The *Slc5a6^cKO^* mice had a lower heart rate, consistent with the known inverse correlation between heart rate and PR interval.^(41)^ CMR imaging revealed that mutant mice had an EF of ∼28.8% at 20 weeks. Patients presenting with an EF of less than 40% are diagnosed with heart failure. ^(42, 43)^ This correlated well with reactivation of the fetal gene programme^(44–46)^ and increased fibrosis, which causes myocardial stiffness and is a characteristic feature of DCM.^(47, 48)^ In human DCM there is an initial adaptive response whereby an increase in collagen fibres boosts growth of the myocardium to enhance contractile force,^(49)^ and this correlates with our proteomic data, showing an increase in collagen-related pathways in mutant mice at 8 weeks. However, over time, this process becomes pathological resulting in impaired cardiomyocyte contractility, causing stiffening of the myocardium, reducing SV and EF until heart failure occurs.

*SLC5A6* facilitates the uptake of dietary biotin and pantothenic acid across the cell membrane. Biotin is an essential cofactor for several carboxylases: acetyl-carboxylase (ACC), PCC, MCC and PC. Pantothenic acid is required for the synthesis of CoA.^(26, 27)^ The metabolomic analysis of the *Slc5a6^cKO^* mouse hearts confirmed deficiencies in both biotin and pantothenic acid metabolism.

Due to the deficiency of biotin, MCC activity, which is required for leucine metabolism, is reduced and the breakdown of 3-methylcrotonyl-CoA occurs via an alternative pathway, leading to the accumulation of C5OH and 3-HIA; metabolites that were elevated in both *Slc5a6^cKO^* hearts and in the patients.^(24, 50, 51)^ Similarly, PCC catalyses the conversion of propionyl-CoA to methylmalonyl-CoA, which is also a crucial step in the metabolism of BCAA (including isoleucine and valine). PCC deficiency leads to the accumulation of 2-methylcitrate,^(52)^ a metabolite that was elevated in the *Slc5a6^cKO^*mouse hearts. Furthermore, three downregulated pathways identified by proteomic analysis are linked to BCAA catabolism. Additionally, pantothenic acid is converted through a series of steps into CoA. Pantothenic acid itself and two of its downstream derivatives, Phosphopantothenic acid and Phosphopantethein were all decreased in the *Slc5a6^cKO^*mouse hearts.

These findings confirm that cardiomyocyte-specific *Slc5a6* deletion disrupts vitamin uptake and mirrors the metabolic abnormalities observed in patients with *SLC5A6* mutations, as well as those with PCC and MCC mutations. Reduced vitamin transport into the heart, therefore, leads to elevated BCAA levels and reduced expression of BCAA catabolism enzymes which have been observed in DCM patients.^(53)^ Enhanced BCAA oxidation in a transverse aortic constriction mouse model improved cardiac function,^(53)^ which supports our hypothesis that biotin and pantothenic acid supplementation could serve as an alternative therapeutic strategy to improve cardiac function by preventing a block in BCAA catabolism.

Of the patients with *SLC5A6* mutations reported in the literature, 64% received different combinations of vitamin supplementation at different stages of disease presentation, resulting in clinical improvements.^(11–15, 17–23)^ In Family 1 reported here, for patient II-1 no vitamin treatment was given and she died from heart failure before the opportunity of a heart transplant arose. Her sibling, patient II-2, vitamin treatment was not initiated until 14 years of age, 7 years after the heart transplant. Modest improvement in development and growth was seen over the past six years, associated with much improved control of her epilepsy. In Family 2,^(18)^ biotin supplementation was initiated 3 weeks after admission for heart failure, and then pantothenic acid was added at day 64, leading progressively to the reversal of the cardiac phenotype and maintained normal cardiac function.

To evaluate whether vitamin supplementation could affect disease progression in our *Slc5a6^cKO^* mouse model, dietary biotin and pantothenic acid were given to the parents before conception and to the pups after weaning. The vitamins were administered at high but safe pharmacological concentrations comparable to doses given to patients, ^(28)^ thereby facilitating cellular uptake of vitamins via passive diffusion^(11)^ and bypassing the need for a functional vitamin symporter in *Slc5a6^cKO^*mice. Interestingly, in the clinical setting, patient II-5 from Family 2 showed no response to an initial 10mg dose of biotin but experienced rapid clinical improvement following an increase to 15mg, supporting the notion that diffusion-mediated uptake can be therapeutically relevant at higher concentrations. Lipoic acid was not supplemented in our mouse model, as it is synthesized endogenously in the mitochondrial matrix and does not require dietary intake in mammals, unlike biotin and pantothenic acid. Remarkedly, vitamin-supplemented mice were phenotypically normal and showed no evidence of cardiac dysfunction or fibrosis and survived until they were electively culled at the end of the study period (40 weeks). These findings underscore the potential of early high-dose vitamin therapy to mitigate cardiac pathology in the absence of *SLC5A6* function. Future research should explore the effects of initiating treatment at different developmental stages, as this could inform optimal therapeutic windows in humans. Importantly, our findings suggest that incorporating *SLC5A6* into prenatal screening could facilitate early diagnosis and timely intervention, potentially improving long-term clinical outcomes.

Previously, there had not been a focused analysis of the heart in patients with SLC5A6 mutations. The results presented here provide a detailed overview of the impact of vitamin loss specifically within the heart on cardiac function and highlights that the underlying initiation of clinical symptoms may be due to a defect in energy metabolism from mitochondrial abnormalities. Imbalances in energy generation or energy use are thought to be key factors in the pathogenesis of DCM^(43)^, which ultimately leads to heart failure.^(54)^ Given the critical role of mitochondria in energy metabolism, alterations in mitochondrial dynamics and function have been associated with cardiovascular diseases and heart failure.^(4)^ However, it is unknown whether the metabolic energetic dysfunction is a cause or a consequence of disease in DCM. Given the link between the vitamins transported by SMVT and mitochondrial energy metabolism, the morphology of mitochondria was analysed as changes in mitochondrial structure can be a pathological factor in disease progression^(29)^ and alterations are known to be associated with functional deficiencies.^(30)^ In cardiomyocytes, the mitochondria are arranged in a highly ordered arrangement, forming rows parallel to the myofibrils,^(55)^ but this organised assembly is lost in *Slc5a6^cKO^* hearts. The mitochondrial complexity index (MCI), an indicator of mitochondrial morphology,^(29)^ was considerably increased in the *Slc5a6^cKO^* hearts by 20 weeks, with increased branching and abnormal donut-shaped mitochondria which are indicative of cellular stress.^(56)^ Mitochondria morphology could be an early indicator of disease onset as smaller yet significant changes in MCI were observed at 8 weeks, alongside the downregulation of key energy metabolism pathways, prior to the onset of an overt DCM phenotype. Interestingly, vitamin supplementation prevented the long-term alterations in mitochondrial morphology in the *Slc5a6^cKO^*hearts. This was also reflected in proteomic data which showed there was no longer a downregulation in mitochondrial metabolism pathways.

Our findings underscore the critical importance of early diagnosis and intervention in *SLC5A6*-related DCM. Vitamin supplementation, when administered early, not only prevents the onset of mitochondrial abnormalities but also halts the progression of DCM, offering a promising therapeutic approach for patients with metabolic-related cardiac diseases. The *Slc5a6^cKO^*mouse model has proven to be an invaluable tool for unravelling the complex relationship between vitamin transport, mitochondrial energy metabolism, and cardiac dysfunction. These results pave the way for novel strategies focused on enhancing mitochondrial function, such as vitamin supplementation, which could preserve cardiac function by boosting energy metabolism^(57)^ potentially transforming the treatment landscape for DCM and other energy metabolism-related heart conditions.

## Supplementary Data

## Acknowledgements

We would like to thank the Newcastle University Electron Microscopy Unit (grant numbers BB/M012093/1 (Gatan 3View system) and BB/R013942/1 (Hitachi TEM) and in particular Ross Laws. The proteomics data was acquired, processed and analysed at the NUPPA core facility, by both Pawel Palmowski and Andrew Porter. We thank Dr Amy Vincent for her invaluable knowledge and insight regarding mitochondrial structure and function. Guido Bommer for his technical help with the LC-MS-qTOF ion-pairing Agilent analysis. Thanks to Prof Judith Goodship for her work on investigating families with heart disorders which led to the discovery of the mutation. Thanks to MRes students Rania Taufiq, Calum Earl, and Brian Satria for their contribution towards the 3D reconstructions of the mitochondria.

## Data availability statement

The proteomic data underlying this article are available in MassIVE Repository at <u>Welcome to MassIVE</u> User: MSV000097747_reviewer. Password: PhillipsH. ftp://MSV000097747@massive-ftp.ucsd.edu

