## Supplementary Material for "A *Slc5a6* Deficient Mouse Model Reveals a Metabolically Driven Dilated Cardiomyopathy with Therapeutic Potential for Vitamin-Based Intervention"

### Methods and Materials

#### Animals

*Slc5a6* transgenic mice (*Slc5a*$^{Tm1a(EUCOMM)Wtsi}$) were obtained from the EUCOMM/IMPC consortium (Figure S2A). The construct included a Neomycin (neo)/LacZ cassette flanked by FRT and loxP sites, inserted between exon 6 and 7. Additional loxP sites were inserted upstream (5') of exon 7 and downstream (3') of exon 10, to enable targeted removal of these four exons within *Slc5a6* using *Cre* recombinase. *Slc5a6*$^{+/Tm1a}$ mice were mated to generate homozygous *Slc5a6*$^{Tm1a/Tm1a}$ mice. In parallel, the neo/LacZ cassette was removed by mating with FLP recombinase mice[(1)] to generate *Slc5a6*$^{Tm1c}$ mice. Homozygous *Slc5a6*$^{Tm1c/Tm1c}$ were further mated with *TnT-Cre* mice[(2)] to delete the function of *Slc5a6* specifically within cardiomyocytes from embryonic day (E)7.5, to produce *Slc5a6* cardiac conditional knockout *Slc5a6*$^{Tm1c/Tm1c}$*;TnT-Cre*$^{+/-}$ (referred to as *Slc5a6*$^{cKO}$) mice (Figure S2A).

Mice were genotyped by standard PCR using primers for *Slc5a6* and *Cre*.

| **Primer** | **Sequence** |
|---|---|
| **Genotyping** | |
| *Slc5a6LoxP2F* | CCTGTCTCTTGAGAAGCCA |
| *Slc5a6LoxP2R* | GGCTGTTGGGAAGCTGAGAT |
| *CreF* | GCATAACCAGTGAAACAGCATTGCTG |
| *CreR* | GGACATGTTCAGGGATCGCCAGGCG |

For vitamin supplementation, breeding cages were provided with vitamin-supplemented mouse diet (TD. 200233 custom diet, ENVIGO) containing 120mg/kg biotin and 2055mg/kg and pantothenic acid, in food pellets. Additionally, 1mM biotin and 1mM pantothenic acid was supplied through drinking water. Vitamin supplementation was initiated prior to conception and maintained throughout the offspring's life.

All studies involving animals were performed in accordance with the UK Home Office Animals (Scientific Procedures) Act 1986 and all experiments were approved by Newcastle University Animal Welfare and Ethical Review Body.

**Reverse Transcription PCR (RT-PCR) and Quantitative real-time PCR (qRT-PCR)**

Total RNA was isolated from mouse tissue using the ReliaPrep RNA Miniprep kit (Promega). cDNA was synthesised using the high-capacity cDNA reverse transcription kit (Thermo Fisher Scientific). For validation of the *Slc5a6*$^{cKO}$ mouse model, RT-PCR was used to confirm the removal of exons 7-10 specifically in the heart only.

qPCR was performed using SYBR green master mix (Thermo Fisher Scientific, in triplicate on a QuantStudio 7 Real-Time PCR System). Primers were designed for targets: CI: *Ndufb8*, CII: *Sdhb*, CIII: *Uqcrc2*, CIV: *Mtco2* and CV: *Atp5a1*. Data were analysed using the comparative Ct method[(3)] and were normalised to *Gapdh* and *β-actin* housekeeping genes and, data shown as the fold change relative to control samples.

| **Primer** | **Sequence** |
|---|---|
| **RT-PCR** | |
| *Slc5a6Ex6F* | ACCATCCTTGGCCCTCAATG |
| *Slc5a6Ex11R* | AGAGACAGGCAACGAAGAGC |
| **qPCR** | |
| *Ndufb8F* | TGGGGTGAACCGATACACTG |
| *Ndufb8R* | GTAAGGGTACTGCTTCGGACC |
| *SdhbF* | CGTTCTCGGCAGAGTCGG |
| *SdhbR* | ACCATAGGTCCGCACTTATTCA |
| *Uqcrc2F* | CCGGGTCCTTCTCGAGATTTTA |
| *Uqcrc2R* | CCGATTCTTGACAGAGGAGCA |
| *Mtco2F* | TCAACATGAAACCCCCAGCCA |
| *Mtco2R* | GCGGCTAGCACTGGTAGTGA |
| *Atp5a1F* | GCCATTTTGTGCCAGTCGTC |

| | |
|---|---|
| *Atp5a1R* | CATTTTTGGAGACCAGTCCCG |
| *GapdhF* | TGTGCAGTGCCAGCCTCGTC |
| *GapdhR* | TGACCAGGCGCCCAATACGG |
| *β-actinF* | CTGCTCTTTCCCAGACGAGG |
| *β-actinR* | AAGGCCACTTATCACCAGCC |
| *NppaF* | GAGAGAAAGAAACCAGAGTG |
| *NppaR* | GTCTAGCAGGTTCTTGAAATC |
| *Myh6F* | CTACTCCTCTTCCTGCCTGTTC |
| *Myh6R* | GTCCGTCATTCTGTCACTCAAA |

**Histology**

Hearts were collected following cervical dislocation and cryoprotected in 7.5-15% sucrose solution for two hours before freezing in OCT or fixed in 4% PFA and embedded in paraffin wax. Serial tissue sections (10µm) were cut and stained with Haematoxylin and Eosin, and Picrosirius Red using standard histological protocols and imaged on a Ziess Axio Imager using brightfield.

**Quantification of Fibrosis: Image Processing and Analysis Macro**

An Ilastik[4] machine learning model was generated using sub-maximum resolution images from the dataset. Partial labelling was applied and the model was trained until sufficient accuracy was reached (determined visually). The model was trained to distinguish tissue of interest from all other tissues/background.

The ImageJ macro was designed within FIJI[5] (ImageJ, NIH) for open accessibility. Images of maximum resolution and images with 4x reduced maximum linear resolution were extracted and saved separately. The lower resolution images were processed using the trained Ilastik model, with output 8-bit probability maps for tissue probability were exported and binaries using pre-processing of a Gaussian blur (sigma=10 pixels) and an automated global thresholding (Huang algorithm). Binary tissue maps were used to remove all regions of images not applicable for further analysis, along with recording the area of each image containing tissue of interest for analysis. Maximum resolution images were separated into individual colour channels, with the green channel values subtracted from the red channel values (to allow for extraction of pixels that show a red stain vs a yellow negative stain). Whole images were measured for pixel intensity in the subtracted red channel, using a minimum threshold value 75, only pixels showing a red intensity above this value are included in analysis.

**Protein Expression**

**Immunofluorescence**: The human embryonic and fetal material was provided by the Joint MRC/Wellcome Trust (grant# MR/R006237/1) Human Developmental Biology Resource (www.hdbr.org). Human embryonic sections were stained by immunofluorescence using the primary antibody SLC5A6 (26407-1-AP, Proteintech) and the nuclei were counter stained with DAPI. Images were acquired on an Axioimager (Zeiss).

Mouse heart sections were stained with wheat germ agglutinin (WGA) conjugated to Alexa Fluor 594, to stain the cell membranes. The cell area of cardiomyocytes was measured using Image J analysis software. Per heart section, four regions of interest images were taken per ventricle, with 3 technical repeats.

**Western Blotting:** Protein expression was quantified using Western blot analysis using standard techniques. Briefly, protein was extracted using protein homogenisation buffer (T-PER, Thermo Fisher, protease inhibitors and phosphatase inhibitors). Samples were homogenised with an electric homogenizer for 30 seconds and centrifuged. Protein lysates

(15000 ng) were denatured in NuPAGE sample buffer and were separated in a 4-15% tris-glycine precast gel with 1xTris-glycine running buffer. Protein was transferred to a PDVF membrane. Biotinylation levels of the carboxylases PCC, MCC and PC were measured using a HRP conjugate[6], IRDye 800CW Streptavidin (Licor), and anti-Gapdh (ab22333). Membranes were imaged and quantified using fluorescent detection on the Licor OdessyFC.

**Protein Modelling**

Three-dimensional protein structures of SLC5A6 were predicted using AlphaFold2[7] accessed via ColabFold.[8] The SLC5A6 amino-acid sequence was obtained from the Uniprot database[9] (UNIPROT ID:Q9Y289) and used as the base query sequence for all structure predictions.

To determine the impact of P437L or R253W point mutations on the SLC5A6 structure, residue substitutions were inserted into the SLC5A6 query sequence as well as into the multiple sequence alignment generated by the wildtype SLC5A6 AlphaFold2 structure prediction.[10] The resulting structure with the highest rank was visualised using Pymol (The PyMOL Molecular Graphics System, Version 2.5.5 Schrödinger, LLC, Available from: http://www.pymol.org/pymol).

**Cardiac Function: Electrocardiography (ECG)**

ECG was performed on anaesthetised mice at 8, 10, 14, 20 and 40 weeks using a 3-lead ECG system (PowerLab).[11] ECG was recorded for three minutes using the PowerLab data acquisition system (ML866, ADInstruments, CO) and animal bio amp (FE136, ADInstruments, CO) on channel 1. Analysis was performed on parameters including heart rate (BPM), PR interval (s), QRS interval (s), ST height (Amp) and T wave (Amp) using LabChart software (AD Instruments). An average of the traces in the third minute was produced in LabChart and quantified when possible.

**Cardiac Function: Cardiac magnetic resonance (CMR) imaging**

CMR imaging was performed on anaesthetised mice using a 7.0T horizontal bore Varian micro imaging system with a 12cm microimaging gradient insert (Varian Inc.). Short axis images were acquired and used to determine cardiac LV function by measuring the LV chamber volume in end systole (EDV) and end diastole (ESV) to evaluate stroke volume (µl), cardiac output (µl) and ejection fraction (%) using ImageJ software (NIH).

**Transmission electron microscopy (TEM)**

Pieces of adult heart (2x2x2mm) were collected and fixed in 2% glutaraldehyde and were stained with heavy metal and embedded in 100% resin.[12] Heavy metal stained sections (70nm) were imaged with CM100 TEM (FEI) in a longitudinal orientation to assess mitochondrial morphology and sarcomeric structure.

**Serial Block Face scanning electron microscopy (SBF-SEM) and 3D reconstructions**

Samples were stained with heavy metal and embedded in 100% resin. Resin blocks were sectioned using in-situ Zeiss Sigma scanning electron microscopy and 250 stacked images were acquired per region of interest (ROI, between 2 and 4). Images taken from each region of interest were combined into a stack using ImageJ (2000 x 2000 x 250 pixels) and the contrast was adjusted to visualise mitochondria throughout. Further analysis was performed using Microscopy Image Browser (MIB, University of Helsinki). Each image stack was aligned and individual mitochondria within a cluster along a myofibril was segmented. Segmented mitochondrial clusters were imported into Amira software (Thermo Fisher Scientific) to generate 3D reconstructions. Amira calculated the volume and surface area of each individual mitochondria. These calculations were used to determine the mitochondrial complexity index (MCI).[13] MCI is used as a quantitative measure of morphological complexity and is the 3D equivalent to form factor (branching) and is analogous to sphericity. The MCI is a parameter used to quantify the 3D shape of mitochondria, irrespective of volume. Thus, spherical mitochondria, whether it be small or large, will have a similar MCI value of ~1. Whereas mitochondria that are elongated, branched, fused or fragmented will have varying MCI values from 0 to infinity.

**Metabolomics**

Acylcarnitine analysis of mouse plasma was performed by flow injection analysis tandem mass spectrometry (FIA-MS/MS) using isotope dilution after a butanol-derivatization step, as part of the routine workflow in the Clinical Biochemical Genetics Laboratory in Brussels. Metabolomic analysis of homogenized frozen heart powder was performed on the upper aqueous fraction after liquid-liquid Folch extraction.[14] CoAG measurement ($[M-H]^-$ 1071.1748) was performed using an ion pairing LC-MS-QTOF Agilent 6550 ion funnel analysis.[14] Other metabolites (3HIA $[M-H]^-$ 117.0562, 2MCA $[M-H]^-$ 205.0362, PA $[M-H]^-$ 218.1051, PPA $[M-H]^-$ 298.072 and PP $[M-H]^-$ 357.088) were measured by LC-MS-qTOF Waters Synapt-XS with an electrospray ionization source in negative mode in a reverse phase chromatography method using an Acquity Premier HSS T3 2.1 x 100mm, 1.8 um column.[15]

**Proteomics**

Protein digestion was carried out with S-Trap micro spin columns (Protifi). Protein concentration was determined using BCA protein assay and an equivalent of 50µg was used for digestion. Proteins were reduced with dithiothreitol (DTT) at a final concentration of 20 mM (65°C, 30min). Cysteines were alkylated by incubation with iodoacetamide (40mM final concentration, 30min, room temp. in dark) and then acidified by adding 27.5% Phosphoric acid to a final concentration of 2.5% (v/v). The samples were then loaded onto spin columns in 6 volumes of loading buffer (90% methanol 100mM TEAB pH 8) and centrifuged at 4000xg for 30s. The columns were then washed with loading buffer (three times) and the flow through discarded. Proteins were digested with trypsin (Worthington) in 50mM TEAB pH 8.5, at a ratio of 10:1 protein to trypsin, overnight at 37°C.

Peptides were eluted with three washes of the trap; first 50 µl 50mM TEAB, second 50µl 0.1% formic acid and third 50 µl 50% acetonitrile with 0.1% formic acid. The solution was frozen then dried in a centrifugal concentrator and reconstituted in 0.1% formic acid 2% Acetonitrile.

Data Independent Acquisition (DIA): 1µl of each peptide sample was loaded per LCMS run. Peptides were separated using an UltiMate 3000 RSLCnano HPLC. Samples were first loaded/desalted onto Acclaim PepMap100 C18 LC Column (5 mm Å~ 0.3 mm i.d., 5 µm, 100 Å, Thermo Fisher Scientific) at a flow rate of 10µL min−1 maintained at 45°C and then separated on a 50cm RP-C18 µPAC™ column (PharmaFluidics) using a 60min gradient from 97 % A (0.1% FA in 3% DMSO) and 3% B (0.1% FA in 80% ACN 3% DMSO), to 35% B, at a flow rate of 400nL min−1. The eluent was directed to a Thermo Orbitrap Exploris 480 mass spectrometer via Thermo Scientific µPAC compatible EasySpray emitter at a temperature of 290°C, spray voltage 1600V. The total LCMS run time was 90 min. Orbitrap full scan resolution was 60,000, RF lens 50%, normalised ACG Target 300%, scan range 390-1600m/z. DIA MSMS were acquired with 49 variable m/z windows covering 390-1621m/z, at 30000 resolution, dynamic maximum injection time, with ACG target set to 3000%, and normalized collision energy level of 30%.

The acquired data was analysed in DIA-NN version 1.8[16] against mouse proteome database (Uniprot UP000000589-2022.06.06) combined with common Repository of Adventitious Proteins (cRAP), Fragment m/z: 300-1800, enzyme: Trypsin, allowed missed-cleavages: 2, peptide length: 7-30, precursor m/z 300-1800, precursor charge: 1-4, Fixed modifications: carbamidomethylation(C), Variable modifications:Oxidation(M), Acetylation(N-term).

The DIA-NN text output was reformatted using an in-house R script and further processed using Perseus (version 1.6.15) software package.

Comparative proteomic analysis between *Slc5a6*$^{cKO}$ mutants and controls for each experimental group, based on diet (normal diet or vitamin supplementation), was performed to identify differentially expressed proteins in Perseus v2.0.9.0.[17] Proteins were filtered for significance ($q<0.05$) and for fold change (FC) $\leq$-0.5 and $\geq$0.5. Pathway analysis was performed using Ingenuity Pathway Analysis (IPA).

**Statistical analysis**

To determine the distribution of all data, a normality test (Shapiro-Wilk) was performed. For data, which was not normally distributed, a non-parametric statistical test was applied (Mann-Whitney). Normally distributed data were subject to parametric testing using an unpaired t-test or one-way ANOVA with Bonferroni correction for multiple comparisons. Mendelian inheritance patterns were assessed with a Chi-squared test. For the mitochondria reconstructions, the slopes of the two regression lines (MCI versus volume) were compared in Prism using an Analysis of Covariance (ANCOVA). Differences between the percentages of mitochondria being simple or complex was tested using a 2x2 contingency table and Fisher's Exact Test. All data are presented as mean ± SEM. $p < 0.05$ was considered statistically significant. Analyses were performed using GraphPad Prism software (version 10.4.1).

**Figure S1. Cardiac clinical data for Family 1 and Family 2 and modelling of *SLC5A6$^{P437L}$* mutation**

(A) Chest X ray of patient II-2 in Family 1 showed an enlarged heart. (B) Pedigree of Family 2. The unaffected consanguineous parents were heterozygous for the SLC5A6 mutation (C/T), whereas patient II-5 had DCM and was homozygous (T/T). Child II-2 died at 8 months from heart failure, but no genetic testing had been performed. (C) Patient II-5 of Family 2 was admitted with an ejection fraction (EF; blue line) of 32%, classified as day 0 and given up to 3 cardiac drugs (grey histogram). The EF returned to normal, allowing discontinuation of treatment, but relapsed and required further cardiac drugs and vitamin supplementation from day 22. The cardiac drugs were stopped on day 148 when the EF became stabilised. Vitamin treatment with biotin and pantothenic acid (PA) is ongoing, and the EF continues to be stabilised. (D) 2D model of SMVT. The P437L mutation is located within transmembrane 11. (E) Amino acid 437 is highly conserved across species. (F-J) 3D modelling of the SMVT protein highlighting P437 in the wildtype (F grey model with P437 in green) and mutant (G orange model with P437L in red) protein. The overall position of the intracellular loops was altered (arrow in H). Substitution with leucine retains the hydrogen bond with residue L441 (magenta) but introduces a novel polar bond with V434 (cyan; black arrow in I,J).

**Figure S2. Generation of *Slc5a6* transgenic mouse models**

(A) The *Slc5a6 Tm1a* allele consisted of a neomycin *lacZ* cassette flanked by *frt* sites (blue triangles) and *LoxP* sites (black triangles) flanking exons 7 to 10 of Slc5a6. The conditional Slc5a6$^{F}$ Tm1c allele was achieved through crossing Slc5a6$^{Tm1a}$ mice with the *Flp* recombinase mouse line to remove the neomycin and *lacZ* cassettes. Subsequent crossing with *Tnt-Cre* transgenic mice removed exons 7-10 within cardiomyocytes, leading to the generation of a premature stop codon (red asterisk) in the Slc5a6 protein. (B-F) Homozygous *Slc5a6$^{Tm1a}$* embryos are embryonic lethal and are underrepresented in litters collected from E9.5 to E15.5. (G) RT-PCR showing *Slc5a6* was expressed in the developing heart at E11.5 to P0. (H-K) SLC5A6 antibody staining in the myocardium of the right and left atria and ventricles at CS14 and CS23 with strong membrane staining (white arrowheads). DAPI was used as a counterstain. (L) RT-PCR confirmed the deletion of exons 7-10 (96bp band) only in the heart (H) and not in the limb (L) of E15.5 embryos. (M) *Slc5a6$^{cKO}$* mice were weaned at the expected Mendelian ratios. (N,O) At 20 weeks, there was a significant decrease in overall body weight (control, *n*=10; Slc5a6$^{cKO}$, *n*=14) but there were no differences in heart weight to tibia length ratio (control, *n*=8; *Slc5a6$^{cKO}$*, *n*=6). (P,Q) Cardiac failure is associated with increased *Nppa* expression and decreased *Myh6* expression in mice, as seen in *Slc5a6$^{cKO}$* 20-week hearts (*n*=3). (R,S) No significant difference was seen in *Nppa* and *Myh6* expression in 20-week vitamin-supplemented Slc5a6$^{cKO}$ mice. Data are represented as mean ± SEM. ns, nonsignificant, *$p < 0.05$ by unpaired t-test. Con, Control; cKO, *Slc5a6$^{cKO}$*, ConV, vitamin-supplemented control; cKOV, vitamin-supplemented *Slc5a6$^{cKO}$*. RV, right ventricle; LV, left ventricle; RA, right atria; LA, left atria; PT, pulmonary trunk; Ao, aorta. Scale bars B-E = 1mm. H,J = 200µm, I,K = 20µm.

**Figure S3. Improvement in cardiac conduction following vitamin supplementation**

(A,B) At 8 weeks, no significant changes in ST height were observed, but the T amplitude was significantly decreased in *Slc5a6$^{cKO}$* compared to control mice. No difference was observed in vitamin-supplemented *Slc5a6$^{cKO}$* mice. (ST height: Con *n*=13, cKO *n*=6, ConV *n*=10, cKOV *n*=9; T amplitude: Con *n*=13, cKO *n*=7, ConV *n*=10, cKOV *n*=10). (C-E) At 40 weeks no significant differences were observed in the vitamin-supplemented *Slc5a6*$^{cKO}$ mice, and all the parameters were comparable to control and vitamin-supplemented control mice. (Con *n*=8, ConV *n*=7, cKOV *n*=6). Data are represented as mean ± SEM. ns, nonsignificant, *$p < 0.05$, by one-way ANOVA with Bonferroni correction for multiple comparisons. Con, control; cKO, *Slc5a6*$^{cKO}$; ConV, vitamin-supplemented control; cKOV, vitamin-supplemented *Slc5a6*$^{cKO}$.

**Figure S4. Measurement of protein biotinylation**

Western blots were probed with streptavidin and Gapdh antibodies. (A,B) Western blots of 5-week hearts and livers on a normal diet (*n*=4). (C-E) Western blots from 20-week hearts and livers on a normal diet and 20-week hearts from vitamin-supplemented diets (*n*=3). PCC and MCC comigrate at 75 kDa, and PC at 130 kDa. ACC1/2 (265 kDa) was not visible on these blots.

**Figure S5. Vitamin supplementation prevents DCM**

Hearts were dissected from 20-week (A,B) and 40-week (D,E) vitamin-supplemented mice. Control and *Slc5a6*$^{cKO}$ hearts were comparable with no evidence of dilation. No difference in body weight was observed at 20 weeks (C) or 40 weeks (F; ConV *n*=7, cKOV *n*=6). (G-K) Picrosirius Red staining of heart sections from 40-week hearts, showed there was no significant increase in fibrosis in the vitamin-supplemented *Slc5a6*$^{cKO}$ mutant hearts. Data are represented as mean ± SEM. ns, nonsignificant, by unpaired t-test. RV, right ventricle; LV, left ventricle; RA, right atria; LA, left atria; ConV, vitamin-supplemented control; cKOV, vitamin-supplemented *Slc5a6*$^{cKO}$. Scale bars A,B,D,E,G,I = 1mm, H,J = 100µm.

**Figure S6. Subtle changes in mitochondria morphology seen in 8-week *Slc5a6*$^{cKO}$ mice**

(A-H) Cardiac mitochondria images from normal diet and vitamin-supplemented hearts from 8-week mice (*n*=3). No obvious differences were seen in TEM images (A,B,E,F). Representative 3D reconstructions of mitochondrial clusters are shown (C,D,G,H). The MCI was calculated for each mitochondria and representative individual mitochondria of the minimum (min), first quartile (1Q), median (med), third quartile (3Q) and maximum (max) MCI are shown (total number of mitochondria analysed: Con = 251, cKO =264, ConV =240 and cKOV =261). (I-L) For 8-week mice on a normal diet the MCI of all of the mitochondria in the $Slc5a6^{cKO}$ hearts was significantly increased, but there was no difference in volume. Bivariate analysis of MCI:volume showed a significant difference in the regression line (K) and the cumulative frequency distribution show partial overlap (L). (M-P) Vitamin supplementation reduced the MCI in $Slc5a6^{cKO}$ mice and the complexity of the mitochondria. Data are represented as mean ± SEM. ns = nonsignificant, *p < 0.05, **p < 0.01, by unpaired t-test. Con, control; cKO, $Slc5a6^{cKO}$; ConV, vitamin-supplemented control; cKOV, vitamin-supplemented $Slc5a6^{cKO}$. Scale bars A,B,E,F = 500nm, C,D,G,H = 1µm.

**Figure S7. Further examples of abnormal mitochondrial morphology in 20-week $Slc5a6^{cKO}$ hearts**

(A-D) TEM images of a control 20-week heart showing the regular arrangement of clusters of mitochondria (white asterisks), parallel to sarcomeres (white arrows). (E-L) TEM images of $Slc5a6^{cKO}$ mice. The mitochondria do not form organised clusters and are randomly dispersed, with irregular sizes and shapes, and evidence of fragmented and damaged mitochondria (black arrows). (M-P) Examples of the different mitochondrial morphologies observed in the $Slc5a6^{cKO}$ mice. Mitochondria with nanotunnels (M), donut shaped (N), elongated (O) and branched (P) mitochondria. Scale bars A,E,I = 2µm, B,F,J = 1µm, C,G,K = 500nm, D,H,L = 200nm, M-P = 0.3µm.

**Supplementary Table 1. Vitamin Doses**

The amount of vitamins given to the mice was calculated for the mouse dose equivalent to human dose from Nair & Jacob (2016)[1], with the assumption that mice eat 3g of food and drink 4ml of water each day. The mean human dose was based on five published studies.[2-6] LC50 is the concentration that results in 50% mortality (death) in a population.

**Supplementary Table 2. Affected subunits of complexes I to V in the electron transport chain in 8-week *Slc5a6$^{cKO}$* hearts**

Gene expression of the different subunits that make up the five complexes of the electron transport chain was analysed in the proteomic data from 8-week control and *Slc5a6$^{cKO}$* hearts ($q < 0.05$, all fold changes). The majority of the subunits present in the proteomic data were downregulated in the Slc5a6$^{cKO}$ hearts: CI 90.2%, CII 100%, CIII 77.8%, CIV 77%, CV 76.5% (highlighted in green) and the non-significant subunits are highlighted in white. The subunits which were not detected in the proteomic data are highlighted in blue. All proteins were downregulated except one complex I subunit, which was upregulated in *Slc5a6$^{cKO}$* hearts (NDUFS6 highlighted in orange).

Figure S1

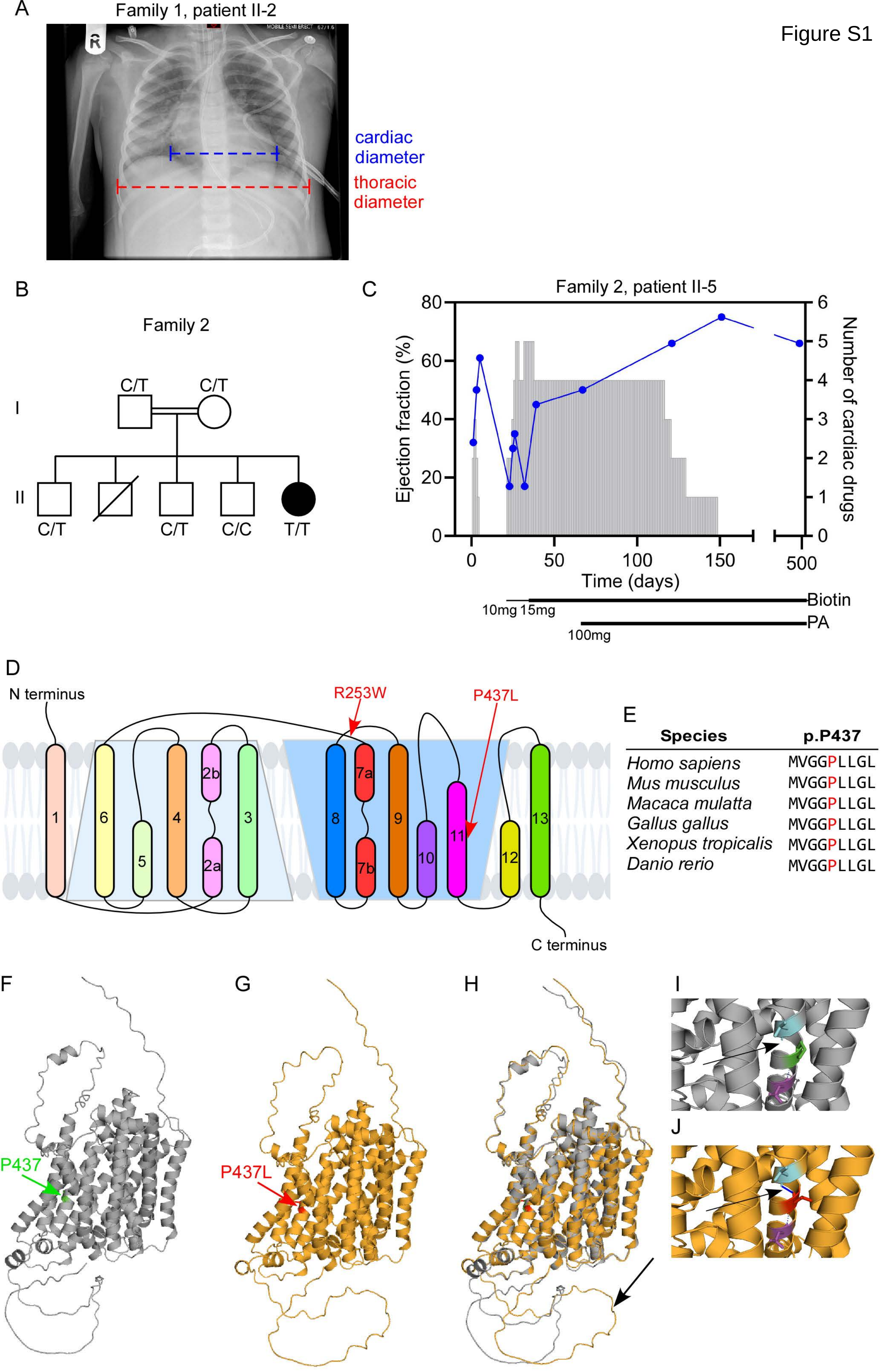

A
Family 1, patient II-2
cardiac diameter
thoracic diameter
B
Family 2
C/T
C/T
I
II
C/T
C/T
C/C
T/T
C
Family 2, patient II-5
Ejection fraction (%)
Number of cardiac drugs
Time (days)
Biotin
PA
10mg 15mg
100mg
D
N terminus
R253W
P437L
C terminus
E
Species
p.P437
Homo sapiens MVGGPLLGL
Mus musculus MVGGPLLGL
Macaca mulatta MVGGPLLGL
Gallus gallus MVGGPLLGL
Xenopus tropicalis MVGGPLLGL
Danio rerio MVGGPLLGL
F
P437
G
P437L
H
I
J

Figure S2

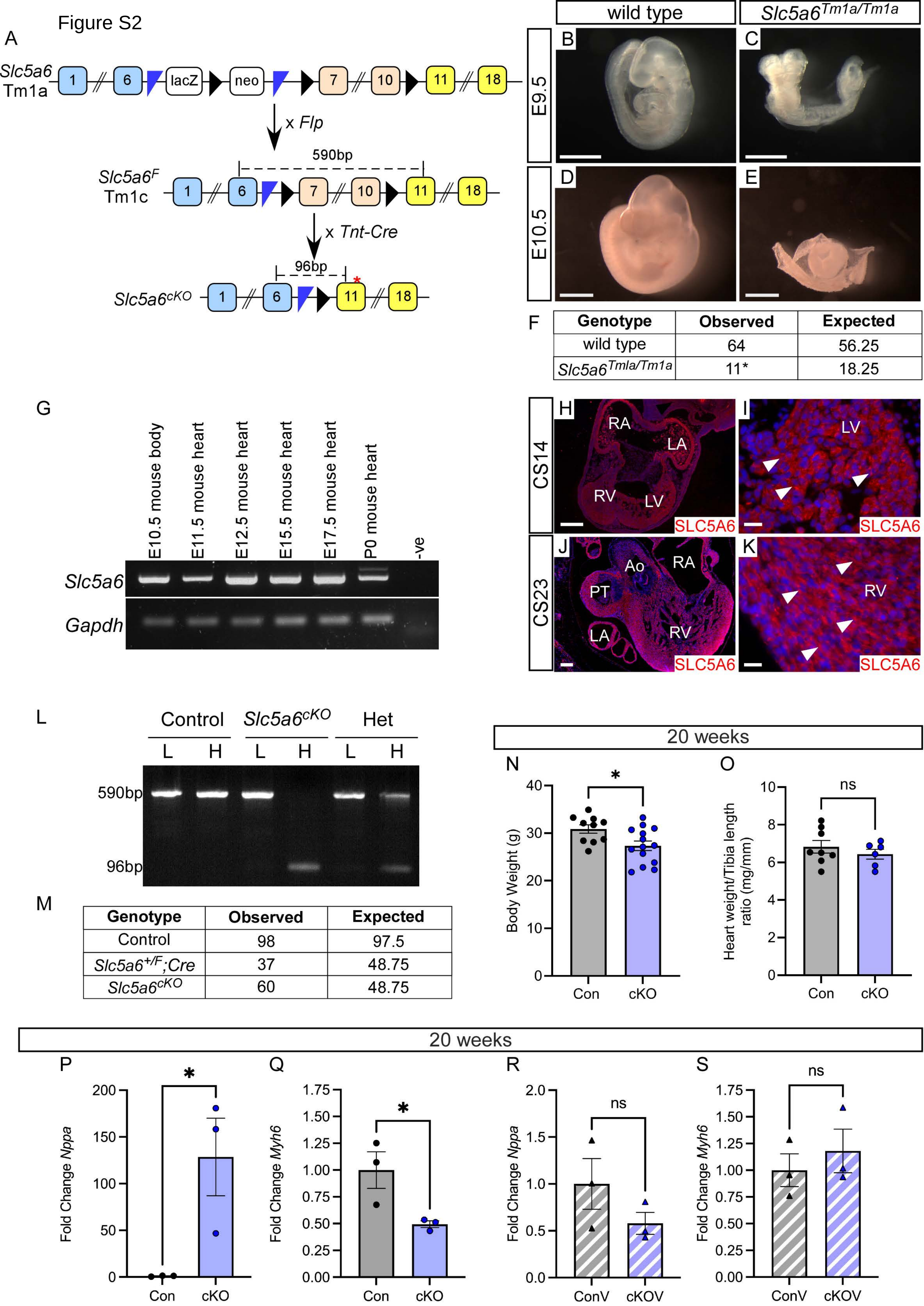

F

| Genotype | Observed | Expected |
|---|---|---|
| wild type | 64 | 56.25 |
| $Slc5a6^{Tmla/Tm1a}$ | 11* | 18.25 |

M

| Genotype | Observed | Expected |
|---|---|---|
| Control | 98 | 97.5 |
| $Slc5a6^{+/F}$;Cre | 37 | 48.75 |
| $Slc5a6^{cKO}$ | 60 | 48.75 |

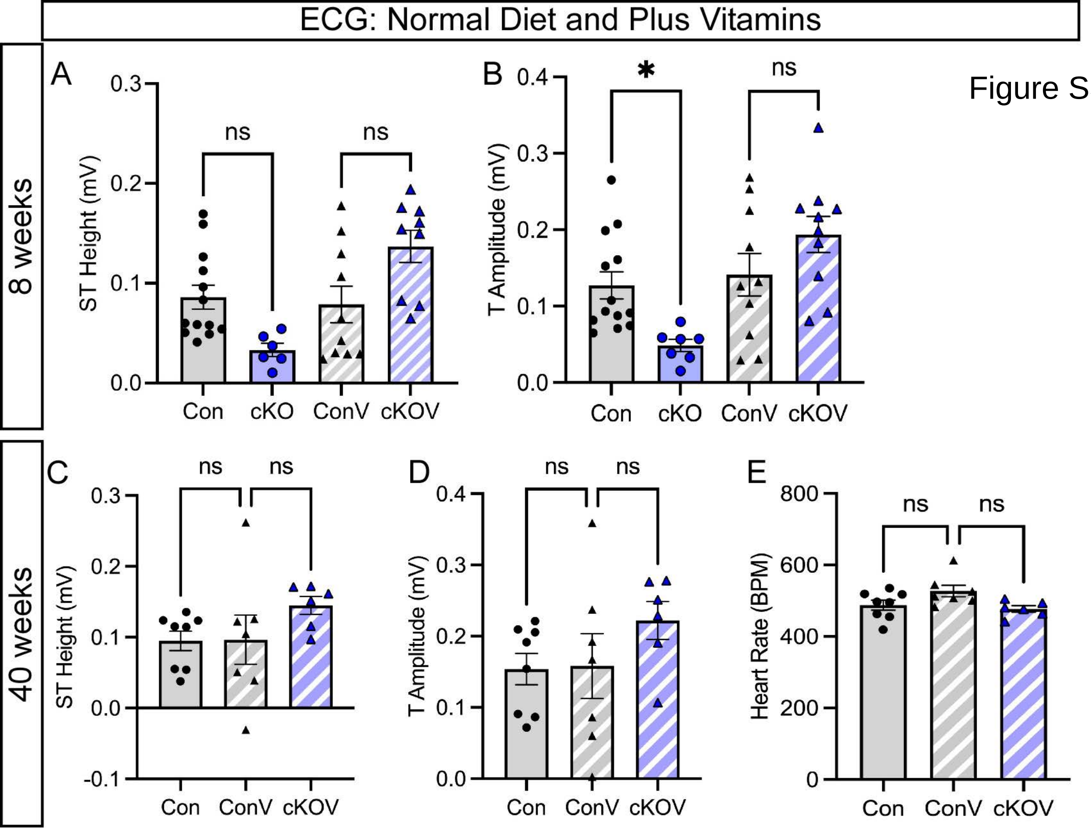

ECG: Normal Diet and Plus Vitamins
8 weeks
40 weeks
A
B
C
D
E
ST Height (mV)
T Amplitude (mV)
Heart Rate (BPM)
ns
*
Con
cKO
ConV
cKOV
0.0
0.1
0.2
0.3
0.4
-0.1
0
200
400
600
800

Figure S3

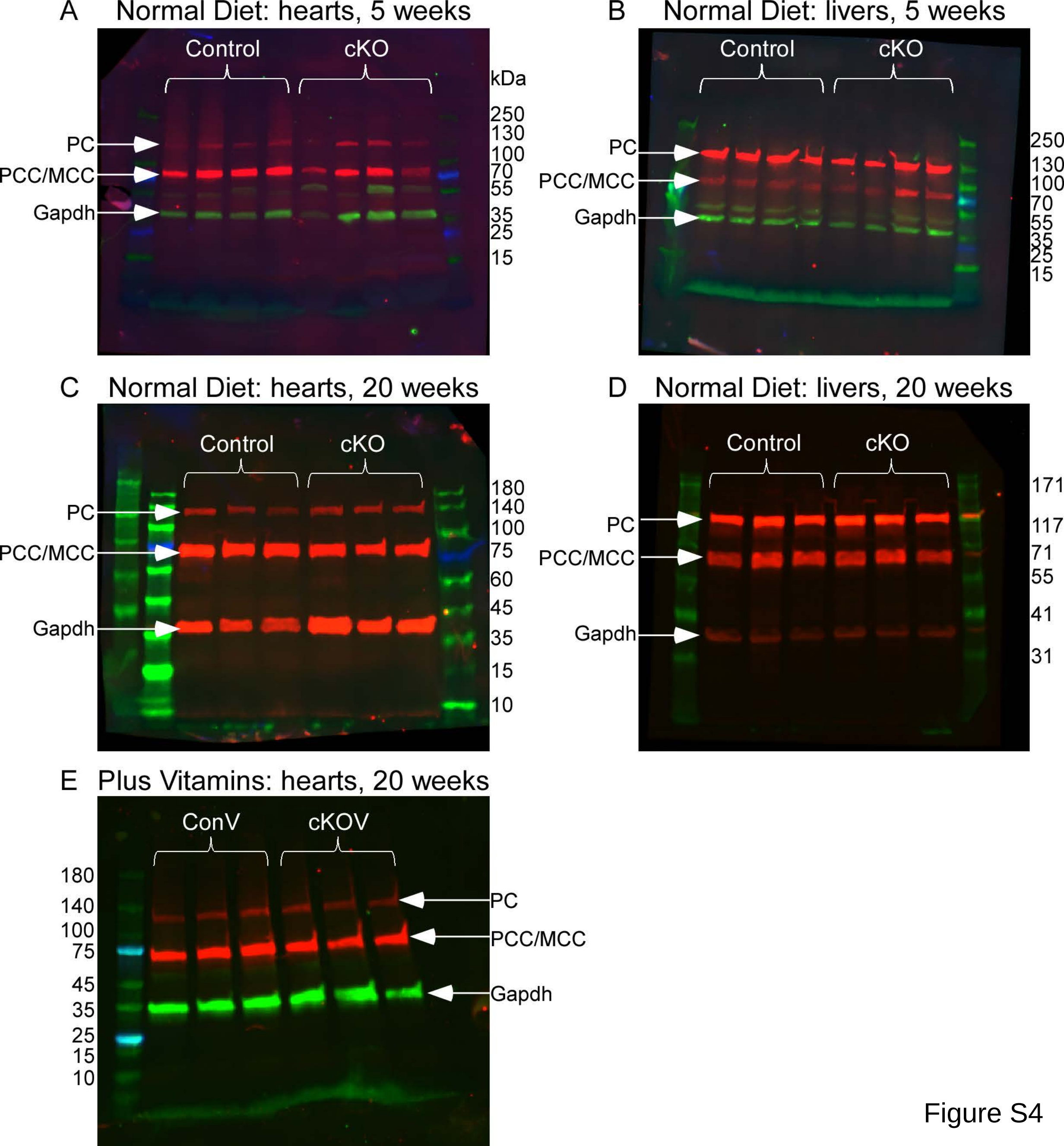
A Normal Diet: hearts, 5 weeks
Control
cKO
kDa
250
130
100
70
55
35
25
15
PC
PCC/MCC
Gapdh
B Normal Diet: livers, 5 weeks
Control
cKO
250
130
100
70
55
35
25
15
PC
PCC/MCC
Gapdh
C Normal Diet: hearts, 20 weeks
Control
cKO
180
140
100
75
60
45
35
15
10
PC
PCC/MCC
Gapdh
D Normal Diet: livers, 20 weeks
Control
cKO
171
117
71
55
41
31
PC
PCC/MCC
Gapdh
E Plus Vitamins: hearts, 20 weeks
ConV
cKOV
180
140
100
75
45
35
25
15
10
PC
PCC/MCC
Gapdh

Figure S4

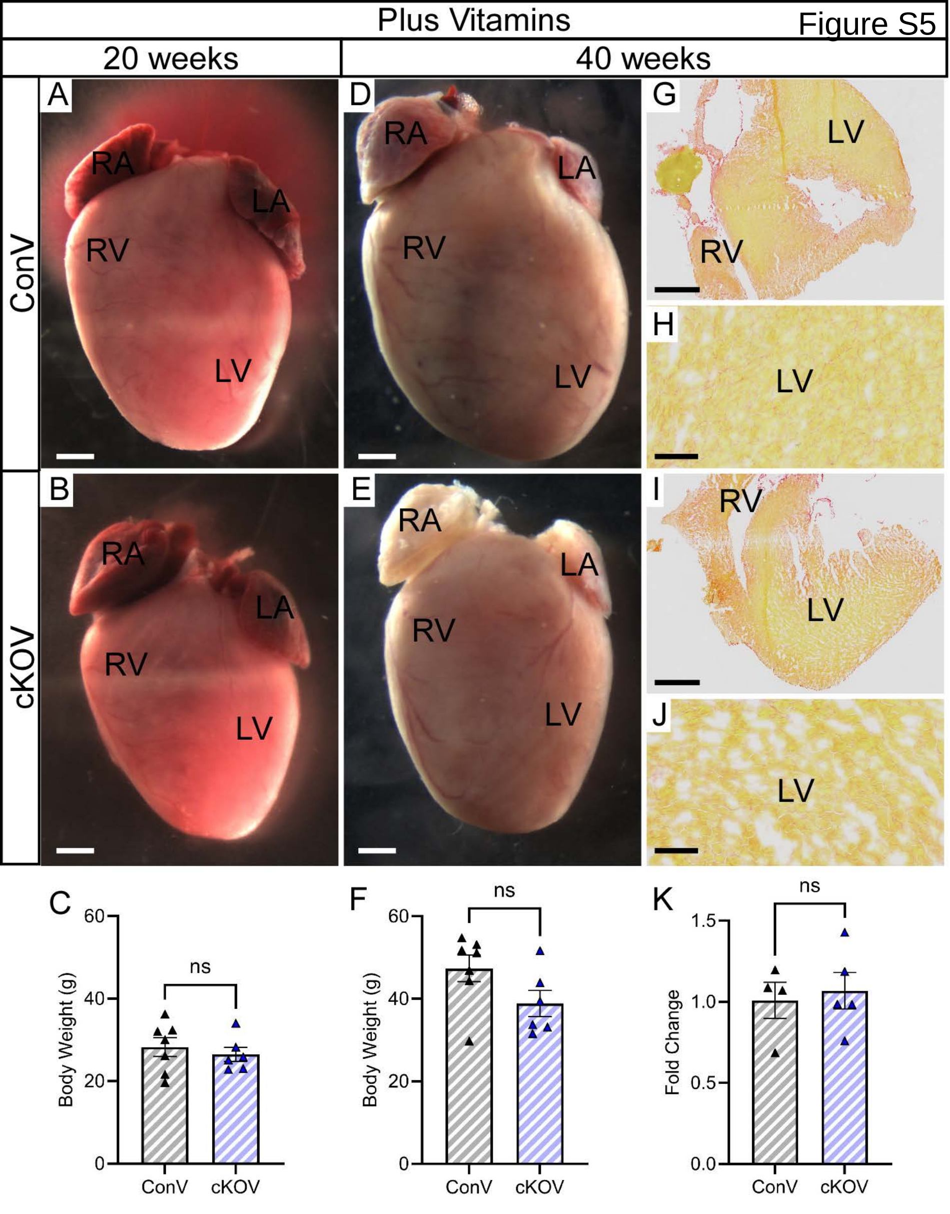
Plus Vitamins
Figure S5
20 weeks
40 weeks
ConV
cKOV
A
RA
LA
RV
LV
B
RA
LA
RV
LV
D
RA
LA
RV
LV
E
RA
LA
RV
LV
G
LV
RV
H
LV
I
RV
LV
J
LV
C
Body Weight (g)
60
40
20
0
ns
ConV
cKOV
F
Body Weight (g)
60
40
20
0
ns
ConV
cKOV
K
Fold Change
1.5
1.0
0.5
0.0
ns
ConV
cKOV

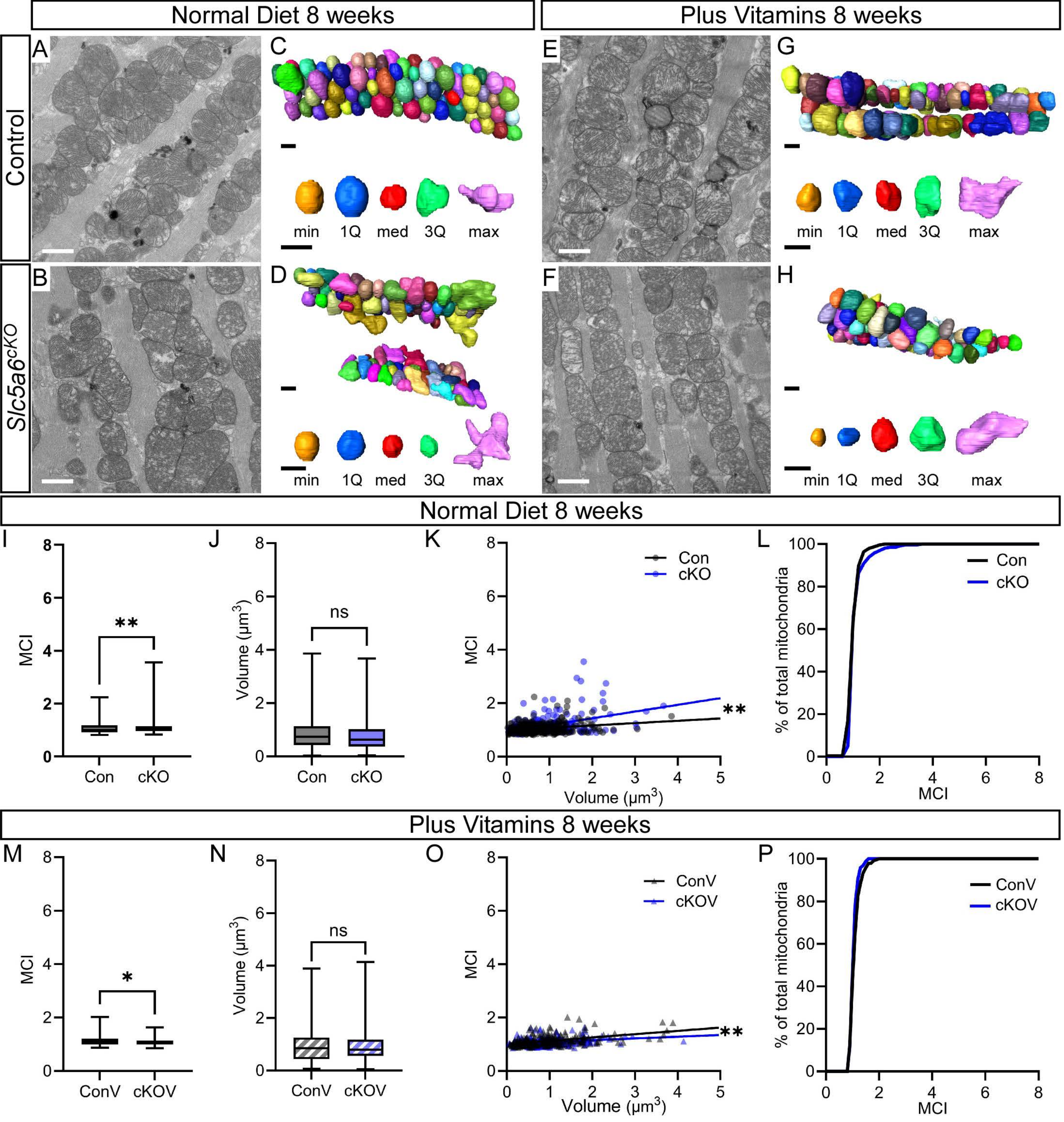

Normal Diet 8 weeks
Plus Vitamins 8 weeks
Control
Slc5a6cKO
A
B
C
D
E
F
G
H
min
1Q
med
3Q
max
Normal Diet 8 weeks
I
J
K
L
MCI
Volume (μm3)
Con
cKO
ns
Volume (μm3)
% of total mitochondria
Plus Vitamins 8 weeks
M
N
O
P
ConV
cKOV

Figure S6

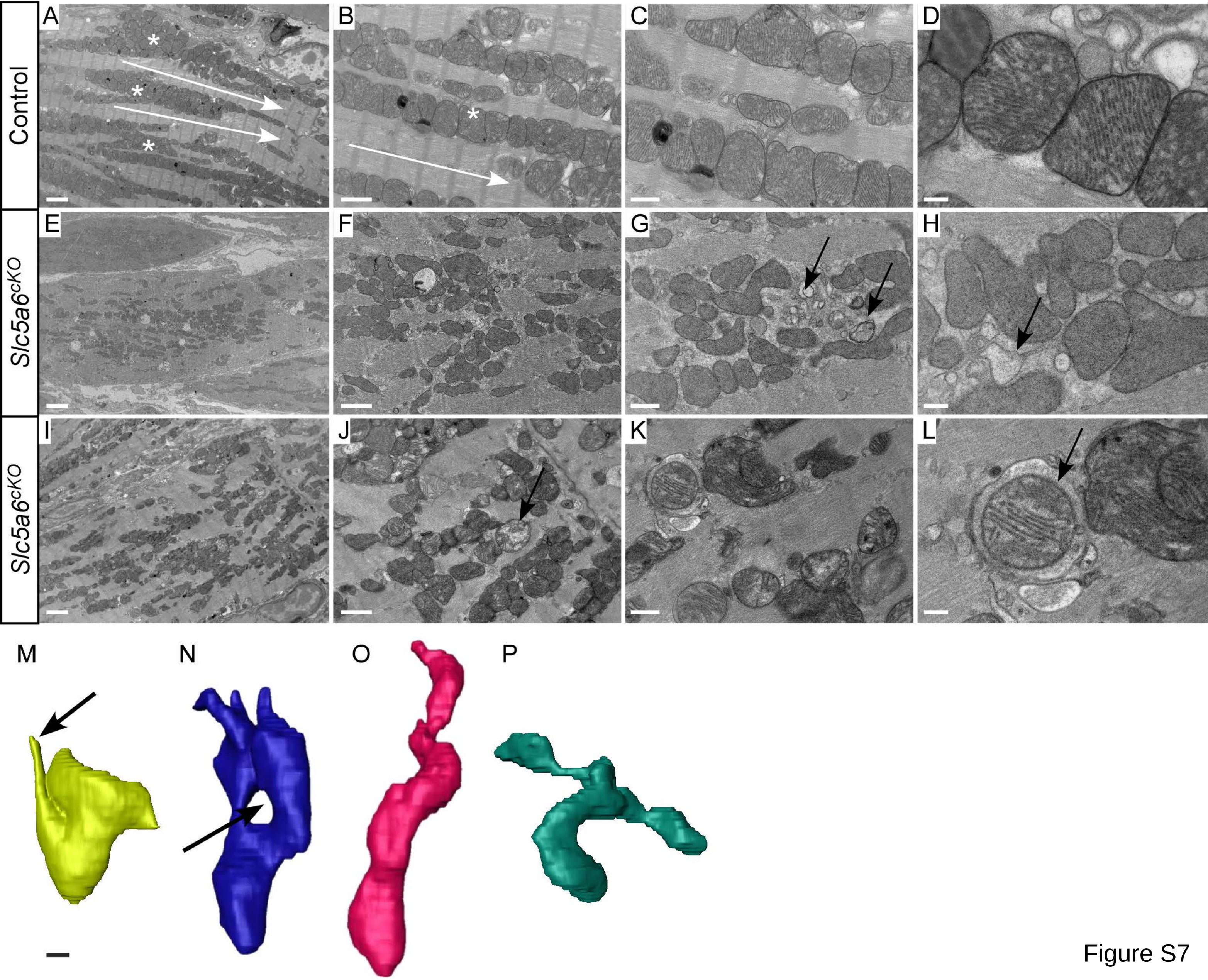

Figure S7

| | $LC_{50}$ (mg/kg) | | Mean dose given (mg/kg/day) | |
|---|---|---|---|---|
| **Vitamin** | **Human** | **Mouse** | **Human** | **Mouse** |
| Biotin | 10,000 | 123,000 | 2.12 | 45.3 |
| Pantothenic acid | 1,443 | 17,749 | 25.9 | 238.7 |

Table S2

| CI | CII | CIII | CIV | CV |
|---|---|---|---|---|
| MT-ND1 | SDHA | UQCRB | COX4I1 | ATP5F1A* |
| MT-ND2 | SDHB* | UQCRQ | COX4I2 | ATP5F1B |
| MT-ND3 | SDHC | UQCRC1 | COX5A | ATP5F1C |
| MT-ND4 | SDHD | UQCRC2* | COX5B | ATP5F1D |
| MT-ND4L | | MT-CYB | COX6A1 | ATP5F1E |
| MT-ND5 | | CYC1 | COX6A2 | ATP5MC1 |
| MT-ND6 | | UQCRFS1 | COX6B1 | ATP5MC2 |
| NDUFS1 | | UQCRH | COX6B2 | ATP5MC3 |
| NDUFS2 | | UQCR10 | COX6C | ATP5ME |
| NDUFS3 | | UQCR11 | COX7A1 | ATP5MF |
| NDUFS7 | | | COX7A2 | ATP5MG |
| NDUFS8 | | | COX7B | ATP5MJ |
| NDUFV1 | | | COX7B2 | ATP5MK |
| NDUFV2 | | | COX7C | MT-ATP6 |
| NDUFAB1 | | | COX8A | MT-ATP8 |
| NDUFA1 | | | COX8C | ATP5PB |
| NDUFA2 | | | MT-CO1 | ATP5PD |
| NDUFA3 | | | MT-CO2* | ATP5PF |
| NDUFA5 | | | MT-CO3 | ATP5PO |
| NDUFA6 | | | | ATP5IF1 |
| NDUFA7 | | | | |
| NDUFA8 | | | | |
| NDUFA9 | | | | |
| NDUFA10 | | | | |
| NDUFA11 | | | | |
| NDUFA12 | | | | |
| NDUFA13 | | | | |
| NDUFB1 | | | | |
| NDUFB2 | | | | |
| NDUFB3 | | | | |
| NDUFB4 | | | | |
| NDUFB5 | | | | |
| NDUFB6 | | | | |
| NDUFB7 | | | | |
| NDUFB8* | | | | |
| NDUFB9 | | | | |
| NDUFB10 | | | | |
| NDUFB11 | | | | |
| NDUFC1 | | | | |
| NDUFC2 | | | | |
| NDUFS4 | | | | |
| NDUFS5 | | | | |
| NDUFS6 | | | | |
| NDUFV3 | | | | |

| | |
|---|---|
| | down in cKO |
| | up in cKO |
| | nonsignificant in cKO |
| | not in proteomic dataset |
| * | subunit in qPCR |
